# Synergistic integration of inducible RNA switches enhances the manipulation of vector expression

**DOI:** 10.64898/2026.02.06.704328

**Authors:** Yueyang Zhang, Yuhan Yang, Zhanzhao Liu, Yue Li, Yizhe Xue, Zihan Zhang, Gonglie Chen, Tian Lu, Yaohua Zhang, Dongyu Zhao, Ke Yang, Lei Miao, Fei Gao, Yuxuan Guo

**Affiliations:** Institute of Cardiovascular Sciences, School of Basic Medical Sciences, Peking University Health Science Center, Beijing, 100191, China; State Key Laboratory of Vascular Homeostasis and Remodeling, Peking University, Beijing, 100191, China; Department of Biomedical informatics, School of Basic Medical Sciences, Peking University Health Science Center, Beijing, 100191, China; Vituner Therapeutics, Nantong, Jiangsu, 226010, China; State Key Laboratory of Natural and Biomimetic Drugs, Beijing Key Laboratory of Molecular Pharmaceutics, School of Pharmaceutical Sciences, Peking University Health Science Center, Beijing, 100191, China; Department of Biochemistry and Molecular Biology, School of Basic Medical Sciences, Peking University Health Science Center, Beijing, 100191, China; Department of Cardiology and Institute of Vascular Medicine, Peking University Third Hospital, Beijing 100191, China; Zhuhai Hengqin SBCVC Xinchuang Equity Investment Management Enterprise (Limited Partnership), Zhuhai, 519000, China; Department of Cardiology, Beijing Anzhen Hospital, Capital Medical University, China; Beijing Key Laboratory of Cardiovascular Receptors Research, Beijing, 100191, China

## Abstract

Precise manipulation of gene expression is pivotal for gene function studies and the optimization of gene therapy. RNA-based gene switches are attractive tools due to their robust tunability by FDA-approved small molecules, the absence of exogenous immunogenic proteins, and the small size for gene delivery vectors such as adeno-associated virus (AAV). However, existing RNA switches only target a single step of gene expression such as transcription or RNA splicing, exhibiting intrinsic limitations in gene regulation. To overcome this issue, this study integrated the aptamer-based polyA regulator (pA), the drug-elicitable alternative splicing module (DreAM) and an engineered translation modulator with conditional upstream open reading frames (uORFs) to construct the DreAM-plus RNA switch. The pA-DreAM concatenation led to 1.5∼5.0-fold and 1.2∼4.4-fold increase of inducible fold changes than pA and DreAM, respectively. The uORF module further enhanced the switching performance by 1.4∼6.3-fold. DreAM-plus-mediated transient transgene expression demonstrated a temporal resolution of about 24 hours and high tissue specificity to liver or heart. Critically, DreAM-plus achieved transient expression of an array of gene editors (SpCas9, SaCas9, Un1Cas12f1, OsCas12f1, AcCas12n, IsDra2 TnpB etc.) that significantly mitigated off-target effects by 1.4∼2.8 folds in plasmids, lentivirus and AAV. In a new mouse model with lipid-nanoparticle-delivered pre-existing immunity, DreAM-plus attenuated AAV-delivered Cas-specific CD8 T cell immune toxicity in the liver and the heart. Therefore, multiple RNA switches could be synergistically integrated to build more sophisticated genetic cassettes for enhanced manipulation of gene expression.

## Introduction

The precise manipulation of gene expression is desirable not only in the study of gene functions but also essential to enhance the efficacy and safety of gene therapy^1,2^. Along the myriad gene expression steps from DNA to protein, inducible switches have been built to control transcription initiation^3–6^, transcription elongation and termination^7^, pre-mRNA splicing^8–11^, mRNA stability^12^, or protein stability and integrity^13–16^. These gene switches usually regulate a single step of gene expression, leaving the other steps uncontrolled and the gene expression with limited tunability. Whether an integrative gene switch that controls multiple layers of gene expression could be built to solve this problem remains unexplored.

Among the diverse gene switching technologies, RNA-based switches are of particular interests. RNA switches are usually free of exogenous proteins, which circumvent their potential immunogenic and pathogenic issues^7,8,10,17^. The absence of protein components also greatly reduces the size of these switches and facilitates gene delivery by vectors with limited payloads, such as adeno-associated virus (AAV). Moreover, accumulative RNA switches have been developed with small-molecule inducers with high safety profiles^7,8,10,11^, increasing their amenability to clinical applications.

To date, RNA switches could be grossly categorized into aptamer- and spliceosome-targeted modules. The aptamer switches are engineered riboswitches that utilize ligand-induced allosteric changes of aptamers to regulate RNA transcription, splicing or stability^18^. Representative RNA switches include the polyA regulator (pA) that controls premature mRNA polyadenylation-transcription termination in response to tetracycline^7^, and the cyclone system that modulates RNA splicing-translation termination by acyclovir^11^. In contrast, the spliceosome-targeted switches take advantage of sequence-dependent small-molecule spliceosome modifiers such as branaplam (LMI070)^8,9^ and risdiplam^10^. This mechanism leads to the development of RNA switches such as Xon^8^ and the drug-elicitable alternative-splicing module (DreAM)^10^.

Among the diverse applications that could benefit from the RNA switches, gene editing is of particular interest due to its broad impact on many different areas. The major side effects of gene editing include off-target effects^19^ and immunotoxicity^20^, which could be reduced by strictly controlling the time and level of editor expression. Electroporation- and lipid nanoparticle (LNP)-delivered gene editing has demonstrated decent safety profiles in clinical studies due to their transient and pulsive expression kinetics^21–23^. However, in other scenarios where electroporation and LNP are not desirable, the gene switch technologies^14,24–26^ are invaluable to overcome gene editing side effects that are associated with plasmids, lentivirus, AAV or other vectors with long-lasting expression^27–30^.

Developing RNA switches for gene editing is challenging for multiple reasons. Firstly, because gene editors could work with a very low concentration, small leakiness of RNA switches is sufficient to compromise their tuning capacity on gene editing^8,9,31^. Secondly, for vectors with limited payloads such as AAV, the demand of delivering the RNA switch together with all gene editing components by a single vector remains unmet^8,16,31,32^. Thirdly, because of the absence of a good model to fully evaluate the immunotoxicity of gene editors^28,33^, the beneficial effect of RNA switches on immune responses remains poorly defined.

To solve these problems, this study developed a new inducible RNA switch by incorporating multiple miniature switching modules such as pA, DreAM and upstream open reading frames that simultaneously control transgene transcription, splicing and translation. In combination with miniature editors such as OsCas12f1, all-in-one AAV vectors with tunable gene editing capacity was successfully constructed. The RNA switch could reduce the expression duration of gene editors and off-target effects. By developing LNP-based animal models with pre-existing immunity against gene editors, this study also demonstrated the plausible application of RNA switches in reducing the immunotoxicity against gene editing.

## Results

### Integration of pA and DreAM enhances drug-inducible gene expression

The Y387 pA regulator utilizes a tetracycline-responsive aptamer to control polyadenylation in the 5’ untranslated region (5’UTR) (Figure 1a) ^7^. To fully understand its working mechanism, a Y387-GFP reporter was assessed in HEK293T cells by reverse-transcription amplicon sequencing (RT-Amp-Seq) and real-time quantitative analysis (RT-qPCR). RT-Amp-Seq detected the splice-in of the aptamer-containing exon in less than 3% mRNA in both solvent- and tetracycline-treated groups (Figure 1b, arrow). RT-qPCR revealed a dramatic increase of vector mRNA by tetracycline treatment (Figure 1c). The Zf2 DreAM module involves a risdiplam-responsive pseudoexon to control start-codon availability (Figure 1d) ^10^. Assessment of a Zf2-GFP reporter by RT-Amp-Seq confirmed risdiplam-induced pseudoexon splice-in (Figure 1e) but no changes in total mRNA were detected by RT-qPCR (Figure 1f).

**Fig. 1.**
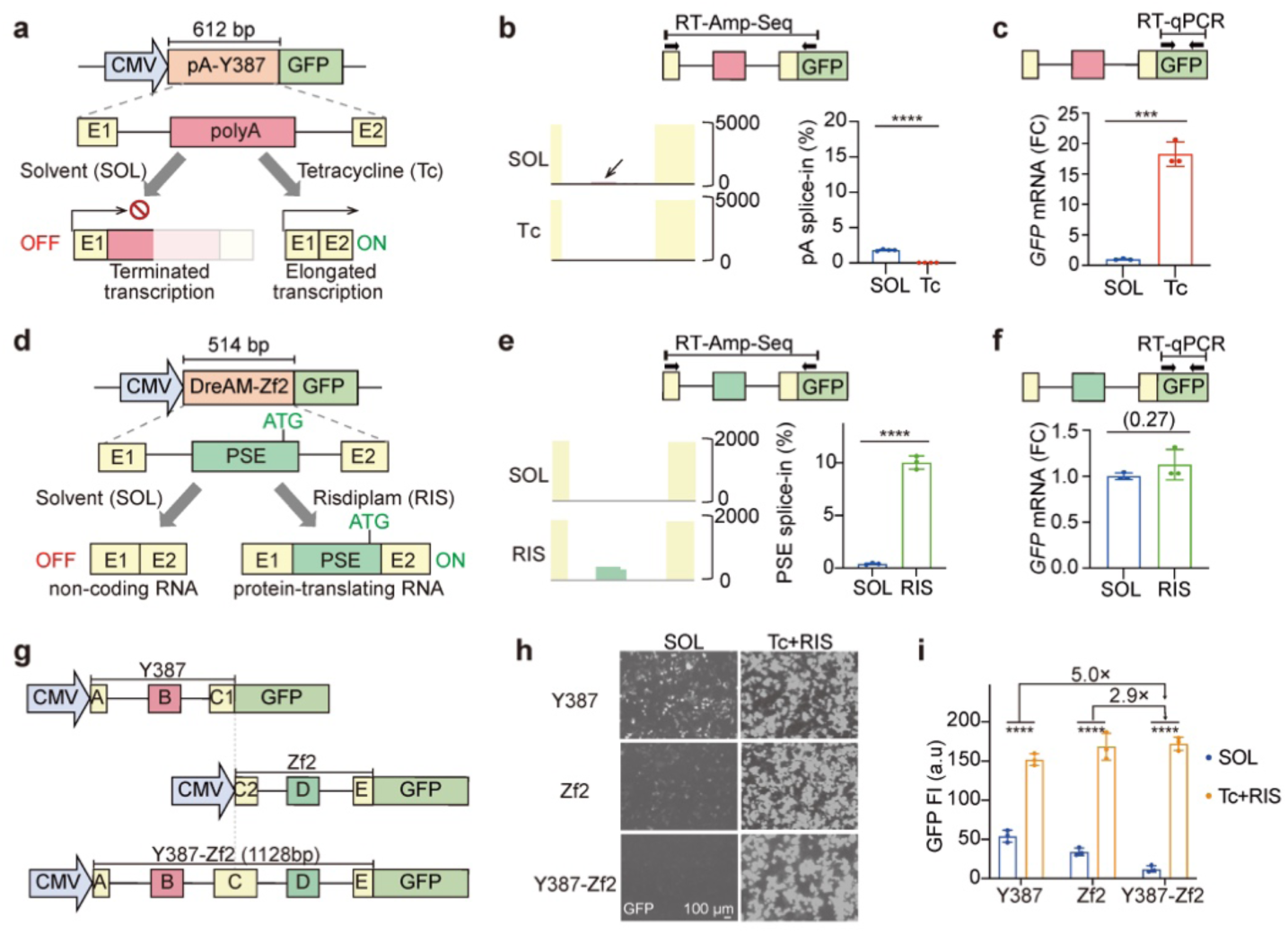
Construction and in vitro characterization of the integrative RNA switch with pA regulator and DreAM. **a,** Schematic illustration of the working mechanism of pA-Y387. E1, exon1; E2, exon2; pA, polyadenylation signal. **b,** Integrated genome viewer (IGV) plots of RT-Amp-Seq reads and quantification of Y387 splicing variants in HEK293T cells treated with 1 μM tetracycline or solvent for 24 hours. The horizontal arrows in the schematic illustration indicate the RT-Amp-Seq primers. The arrowheard in the IGV plot indicates basal splice-in signals without tetracycline. RT, reverse transcription. **c,** Y387-controlled *GFP* mRNA quantification in HEK293T cells. The arrows indicate the quantitative PCR primers. FC, fold change. **d,** Schematic illustration of the working mechanism of DreAM-Zf2. PSE, pseudo-exon. **e,** IGV plots and quantification of Zf2 PSE splice-in rates in HEK293T cells treated with 1 μM risdiplam or solvent for 24 hours. **f,** Zf2-controlled *GFP* mRNA quantification in HEK293T cells. **g,** A diagram showing the structures of Y387, Zf2, and their concatenation to form the Y387-Zf2 fusion element with five exons labeled as A∼E. **h,i,** Representative fluorescence images **(h)** and quantification **(i)** of GFP signals in HEK293T cells treated with 1 μM tetracycline plus 1 μM risdiplam or solvent for 24 hours. a.u, arbitrary units. The fold changes of induced dynamic ranges between the different RNA switches are labeled above. **b,c,e,f** and **i,** n = 3-4 biological repeats. Two-tailed unpaired Student’s t-test: ***P < 0.001, ****P < 0.0001. Non-significant P values are in parentheses.

The orthogonal working mechanisms of Y387 and Zf2 on transcription and splicing suggested an opportunity of integration for better performance. Therefore, Y387 and Zf2 were concatenated to build a new RNA switch containing five exons (Figure 1g). The Y387-Zf2-GFP reporter demonstrated greatly reduced baseline leaky expression than Y387 or Zf2 alone (Figure 1h-i). By combined treatment of tetracycline and risdiplam, the Y387-Zf2 switch exhibited 5.0- or 2.9-fold increases in inducible fold changes than Y387 or Zf2 alone (Figure 1i, ED Figure 1a-b). This enhanced switching performance was also validated by a luciferase reporter in HEK293T, NIH3T3, and Neuro2a cells (ED Figure 1c).

The five exons in the Y387-Zf2 switch were denoted as exon A∼E, which were predicted to variably splice into eight transcripts (Figure 2a). The Y387-Zf2 switch could be induced by tetracycline alone to increase the mRNA level, which was further enhanced by risdiplam (Figure 2b). Because risdiplam did not directly work on Y387 (ED Figure 1d), the impact of risdiplam on the mRNA level of the Y387-Zf2 switch indicated a synergistic effect between transcription and splicing. RT-Amp-Seq analysis of the Y387-Zf2 switch further uncovered mutually exclusive alternative splicing between exon B, C and D (Figure 2c). Risdiplam induced splice-in of exon D, which was originally the pseudoexon in Zf2, could be further elevated by tetracycline (Figure 2c-d), again indicating a synergistic effect between transcription and splicing.

**Fig. 2.**
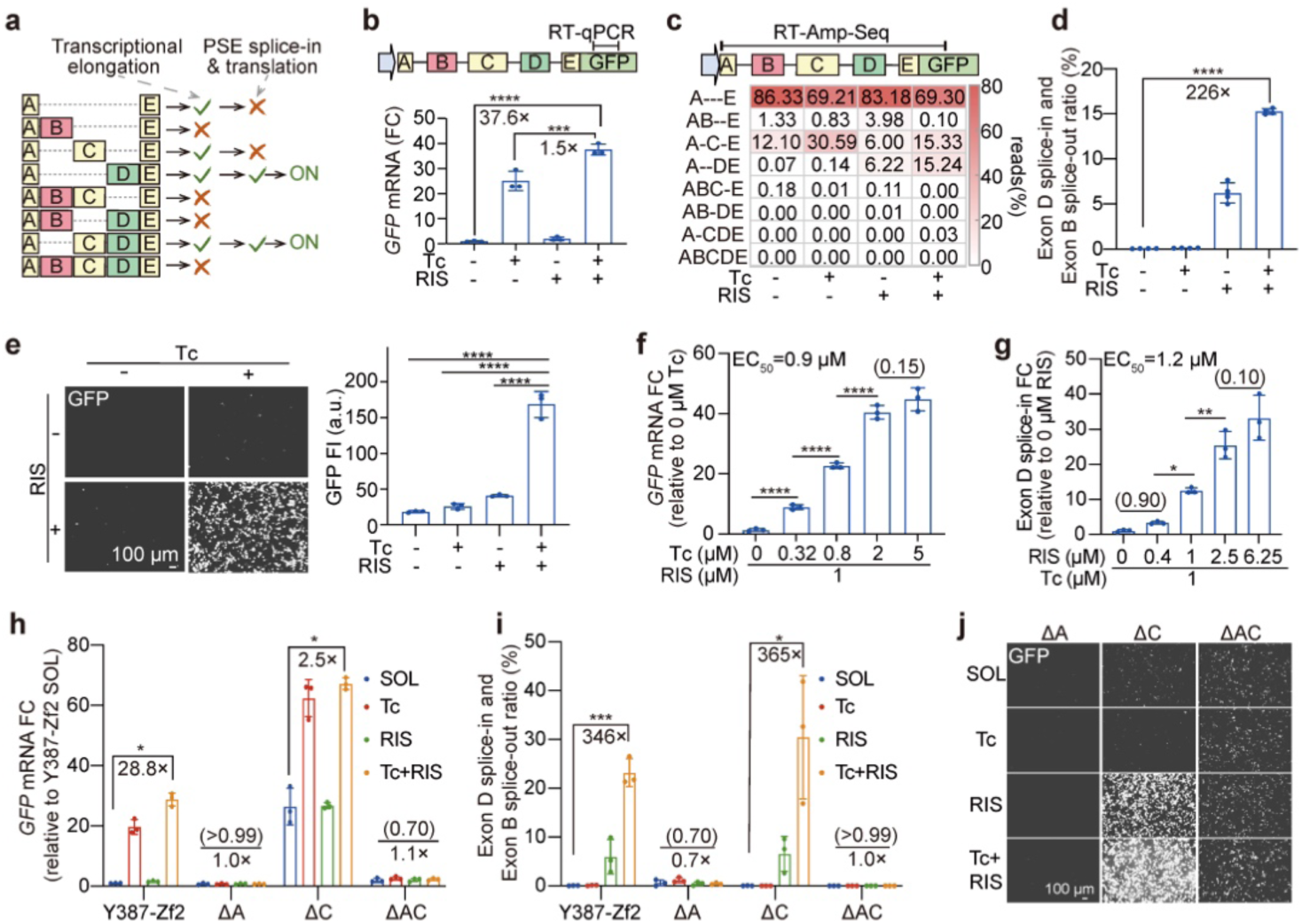
The pA regulator and DreAM integration enables multi-layer regulation of gene expression. **a,** A diagram showing the splicing variants and their respective gene expression status. **b,** *GFP* mRNA quantification of HEK293T cells treated with solvent, 1 μM tetracycline, 1 μM risdiplam or both for 24 hours. **c,d,** Heatmap depicting the relative abundance of different splicing variants **(c)** and quantification of exon D splice-in while exon B splice-out ratios **(d)**. **e,** Representative fluorescence images and quantification of GFP intensity. **f,** Dose response of *GFP* mRNA levels to a gradient of tetracycline concentrations. EC_50_, half-maximal effective concentration. **g,** Dose response of Exon D splice-in ratio to a gradient of risdiplam concentrations. **h-j,** Characterization of exon-deleted switch variants. HEK293T cells transfected with the indicated switch variants were treated for 24 hours before analysis of *GFP* mRNA levels **(h),** exon D splice-in while exon B splice-out ratio **(i)**, and GFP fluorescence **(j)**. ΔA, exon A deletion; ΔC, exon C deletion; ΔAC, exon A and C deletion. **b,d,e-i,** n = 3-4 biological repeats. One-way ANOVA with Tukey’s test: *P < 0.05, **P < 0.01, ***P < 0.001, ****P < 0.0001. Non-significant P values are in parentheses. The fold changes are labeled below P values.

Although tetracycline and risdiplam could individually induce transcription or splicing, both inducers were necessary to induce protein expression by Y387-Zf2 (Figure 2e, ED Figure 2a). Each inducer promoted transgene expression in a dose-dependent manner when the dose of the other inducer was fixed. Tetracycline and risdiplam demonstrated a EC_50_ of 0.8∼0.9 μM and 1.1∼1.2 μM, respectively (Figure 2f-g, ED Figure 2b-c). Exon A deletion abolished the inducibility of Y387-Zf2 (Figure 2h-j) because exon B could not be skipped to suppress 5’UTR polyadenylation (ED Figure 2d). Exon C deletion resulted in higher leaky expression due to the enhanced mRNA level (Figure 2h-j). Therefore, all five exons are essential for its tuning capacity.

### Conditional upstream open reading frames reduce aberrant protein translation

A major caveat of RNA splicing switches that regulate the availability of start codons is the undesired leaky expression of N-terminal truncated proteins^10,17^ (Figure 3a). This issue occurs when the transgene contains alternative start codons adjacent to its 5’ end, allowing protein translation by these downstream open reading frames (dORF) to escape the control by the RNA switch (Figure 3a). The Y387-Zf2 switch greatly reduced the leaky expression of dORF-coding proteins in the Zf2-GFP reporter (ED Figure 1b, red stars), which was more evident when Y387-Zf2 was applied to control gene editors. For example, SaCas9, Un1Cas12f1, TnpBmax and TnpB were founded to express many dORF proteins at the OFF state by Zf2, which could be eliminated by Y387-Zf2 (Figure 3b-d, ED Figure 3a-b, red stars). Amplicon-sequencing analysis further confirmed enhanced inducible fold changes in the deposition of insertions and deletions (indels) under the control of Y387-Zf2 over Y387 or Zf2 alone (Figure 3b-d, ED Figure 3a-b).

**Fig. 3.**
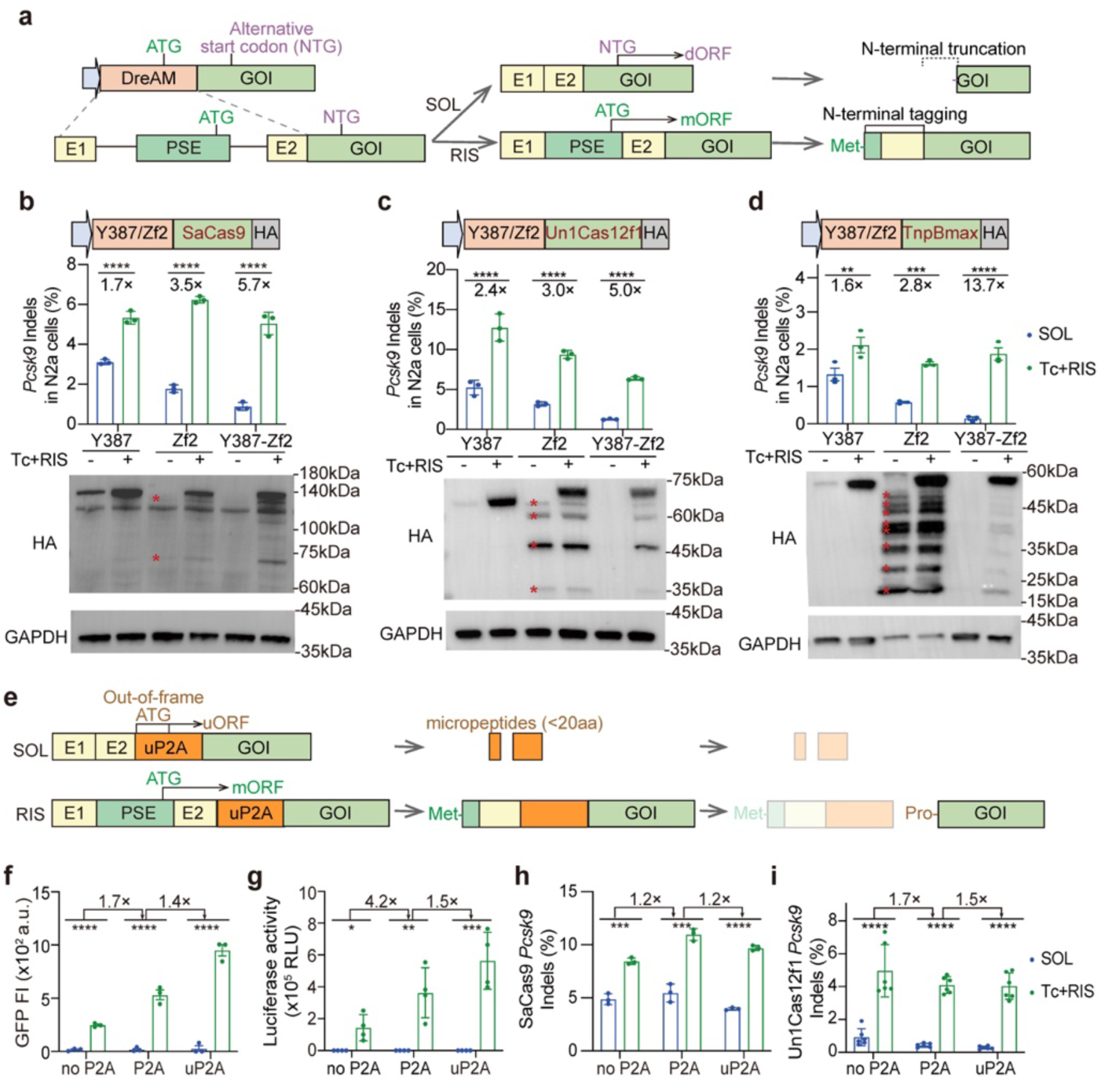
P2A with upstream ORFs reduces aberrant protein translation by DreAM. **a,** A diagram showing the generation of aberrant N-terminal truncated peptides and N-terminal tagged peptide by DreAM. GOI, gene of interest; dORF, downstream open reading frame; mORF, main open reading frame; Met, methionine. **b-d,** Gene editing efficiency and corresponding western blot analysis of SaCas9 **(b),** Un1Cas12f1 **(c)** and TnpBmax **(d)**. The RNA switches were treated for 48 hours with solvent or 0.9 µM tetracycline plus 1.2 µM risdiplam. Red stars denote N-terminal truncated protein bands. Indels, insertions and deletions; N2a, Neuro-2a. n = 3-4 biological repeats. **e,** A diagram showing the mechanism of uP2A for enhanced regulatory capacity. uORF, upstream open reading frame; uP2A, P2A with uORFs; Pro, proline. **f,g,** Quantification of GFP **(f)** and luciferase **(g)** activity controlled by switches with different P2A variants. **h,i,** Gene editing efficiency of SaCas9 **(h)** and Un1Cas12f1 **(i)** controlled by switches with different P2A designs in N2a cells. **f-i,** n=3-6 biological repeats. The fold changes of induced dynamic ranges are labeled above. Two-tailed unpaired Student’s t-test: *P < 0.05, **P < 0.01, ***P < 0.001, ****P < 0.0001.

The RNA splicing switches that regulate the availability of start codons were often coupled with a self-cleavage P2A peptide^9,10^ to remove the extra amino acids that were coded by the RNA switch itself (N-terminal tagging in Figure 3a). To utilize this element for better control of protein translation, P2A was engineered by introducing synonymous mutations to generate out-of-frame start codons and upstream ORFs (uORFs) to reduce the activity of dORFs (Figure 3e, ED Figure 3c). For all tested transgenes including GFP, Luciferase, SaCas9 and Un1Cas12f1, this uORF-containing P2A (uP2A) element enhanced the inducible fold changes by 1.4∼6.3 folds than transgenes lacking P2A (Figure 3f-i). The superior performance of this new RNA switch on gene editing was further validated by additional editors including OsCas12f1 and <u>Ac</u>Cas12n (ED Figure 3d-e). Western blot confirmed the reduction of both dORF leaky expression and N-terminal tagging issues (ED Figure 3f-g).

### DreAM-plus mediates transient and tissue-specific AAV expression in vivo

The new inducible RNA switch that combined Y387, Zf2 with or without uP2A was termed as DreAM-plus (DreAM+), which was next delivered in an AAV9-Luciferase vector to characterize its performance in mice (Figure 4a). A single dose of the inducers (80 mg/kg tetracycline + 50 mg/kg risdiplam) was orally administered at one week after AAV injection before time-lapse luciferase assay was conducted. This experiment detected transient bioluminescent signals that peaked at 12 hours after inducer administration (Figure 4b). This faster turnover rate of DreAM-plus over DreAM^10^ is consistent with the shorter half-life of tetracycline (∼9 h)^34^ than risdiplam (∼50 h)^35^. Over a period of two months, the DreAM-plus-Luciferase reporter was successfully induced for three times to confirm the repeatability in vivo (Figure 4c). Each inducer activated the transgene in a dose-dependent manner when the other inducer was given a fixed dose (Figure 4d-e).

**Fig. 4.**
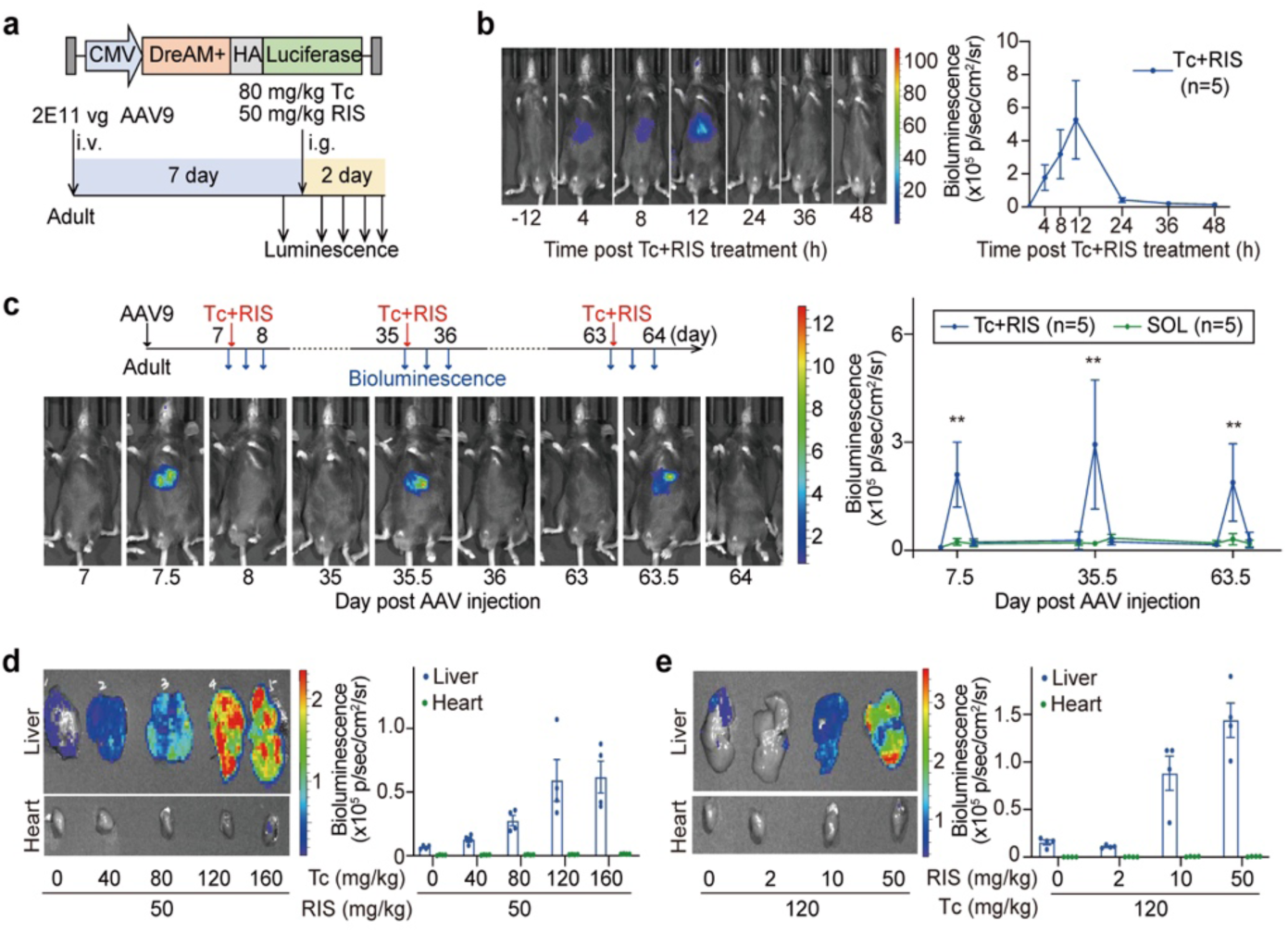
DreAM-plus enables dose-dependent and reversible AAV control in vivo. **a,** Experimental design for assessing the performance of AAV-DreAM-plus in mice. Mice received 1E10 vg/g AAV9 via tail-vein injection, followed by 80 mg/kg tetracycline plus 50 mg/kg risdiplam treatment via intragastric administration. DreAM+, DreAM-plus; i.v., intravenous injection; i.g., intragastric administration. **b,** Representative bioluminescence images and quantification of abdominal signals at indicated time points. n = 5 animals per group. **c,** Bioluminescence analysis for evaluating repeatable induction. Adult mice with AAV9-DreAM+-Luciferase were administered with multiple doses of solvent or inducers. Representative images and abdominal signal quantification were shown. n = 5 animals per group. Two-tailed unpaired Student’s t-test: **P < 0.01. **d,e** Dose-dependent responses to tetracycline **(d)** or risdiplam **(e)** in isolated livers and hearts via i.g. n = 4 animals per group.

Imaging of dissected organs confirmed that the liver was the only organ that exhibited inducible signals in the previous experiment (Figure 4d-e). To figure out the working conditions of DreAM-plus for other organs, Y387, Zf2 and DreAM-plus were compared side-by-side in liver, heart and skeletal muscle via intragastrical (i.g.) or intraperitoneal (i.p.) injections of the inducers (Figure 5a-c). Interestingly, Zf2 responded similarly to i.g. and i.p. injections in all three organs, but Y387 and DreAM-plus favored i.p. over i.g. injection by 5∼35 folds (Figure 5a-c). A single i.p. injection also led to the peak signal at 8-12 hours, which returned to baseline by 24 hours (ED Figure 4a). Dose-dependent Luciferase activity in the heart and skeletal muscle was confirmed upon i.p. injections (ED Figure 4b-c).

**Fig. 5.**
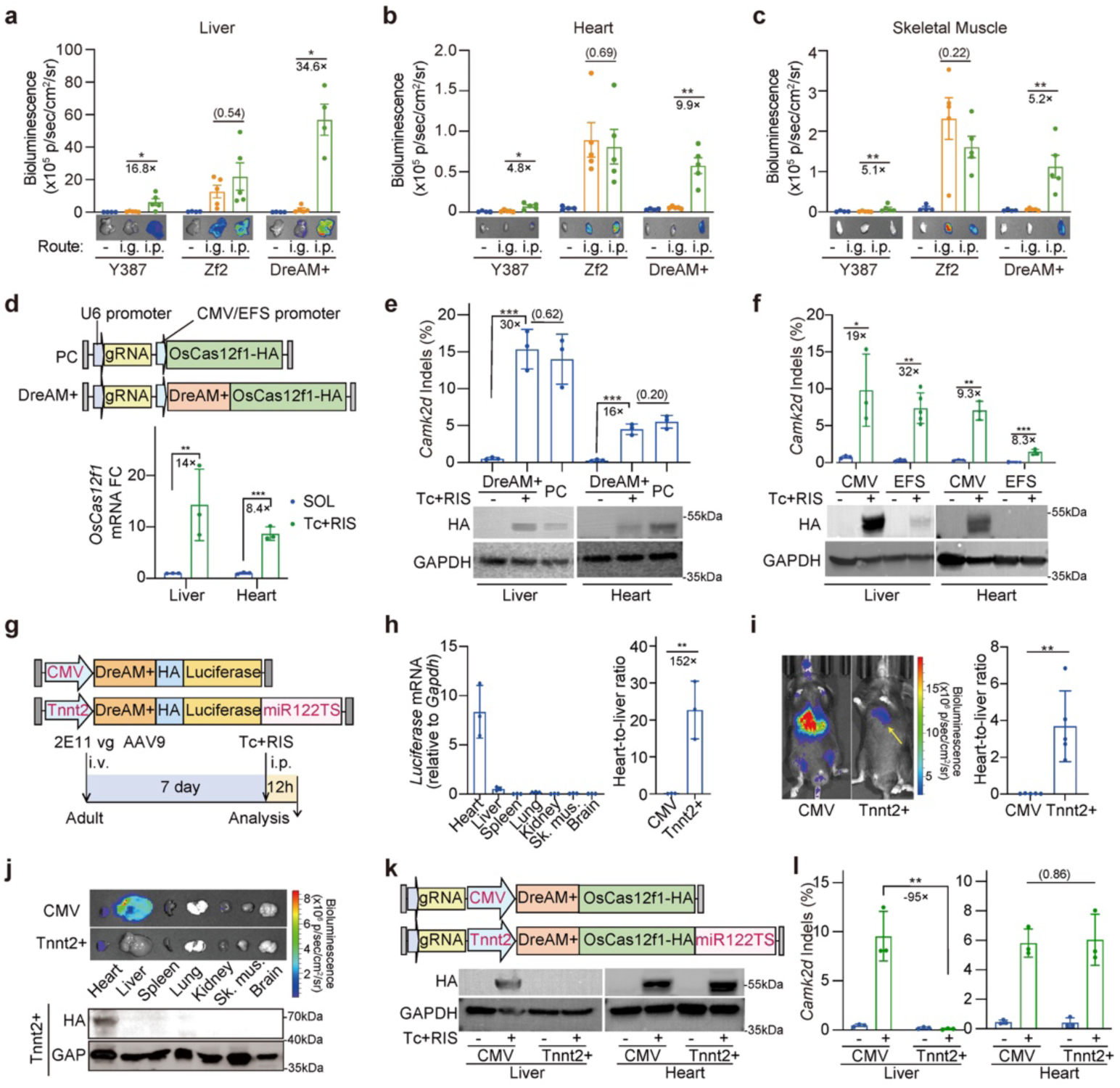
Cardiac-specific AAV regulation by DreAM-plus. **a-c,** Bioluminescence images and quantification of isolated liver **(a)**, heart **(b)** and skeletal muscle (quadriceps femoris) **(c)** via different administration routes. Mice were treated with solvent or 80 mg/kg tetracycline plus 50 mg/kg risdiplam for 12 hours before imaging. n = 4-5 animals per group. **d-f,** Inducible *OsCas12f1* mRNA **(d)**, protein and gene editing **(e-f)** analysis in the liver and the heart. PC, positive control. Ubiquitous promoter strength, CMV>EFS. n = 3 animals per group. **g,** Experimental design to assess the compatibility of DreAM+ with the cardiac specific Tnnt2-miR122TS vector (Tnnt2+) via intraperitoneal injection of the inducers. **h,** *Luciferase* mRNA quantification of the Tnnt2+ vector in different organs (left) and the heart-to-liver mRNA ratio of CMV or Tnnt2+ vectors (right). **i,** Bioluminescence images and quantification of heart-to-liver ratios. The yellow arrow points to the heart region. n = 5 animals per group. **j,** Representative bioluminescence images and western blot analysis of CMV and Tnnt2+ groups. **k,l,** OsCas12f1 western blot and gene editing analysis in the liver and heart. For gene editing analysis, n = 3-5 animals per group. Two-tailed unpaired Student’s t-test: *P < 0.05, **P < 0.01, ***P < 0.001. Non-significant P values are in parentheses.

To construct an all-in-one AAV vector carrying DreAM-plus regulated gene editing, the miniature Un1Cas12f1 (1.6 kb coding sequence)^36^ and OsCas12f1 (1.3 kb coding sequence)^15^ were adopted. Under the control of the CMV promoter, a single i.p. injection of the inducers (80 mg/kg tetracycline + 50 mg/kg risdiplam) resulted in enhanced gene editor expression in both the liver and the heart (ED Figure 4d and Figure 5d-f). Amplicon sequencing confirmed the inducible deposition of indels at the levels that were comparable to canonical AAV-OsCas12f1 treatment (PC in Figure 5d-e). When a weaker ubiquitous promoter (EFS) ^8,9,37,38^ was side-by-side compared to the CMV promoter, both baseline and induced gene editing were reduced while the inducible fold changes were barely compromised (Figure 5f). Notably, the heart always exhibited lower inducible fold changes than the liver because both inducers were preferentially enriched in the liver over the heart^10,39^.

Next, DreAM-plus was incorporated into an AAV9-Tnnt2-miR122TS vector^40^ to achieve cardiac-specific induction (Figure 5g). Tnnt2 (cardiac troponin T) is a cardiomyocyte-specific promoter^41^ and miR122TS (microRNA-122 target sequence) is a robust liver-detargeting element^42^. RT-qPCR, luciferase assay, and western blot confirmed up to 152-fold increase of the heart-to-liver signal ratios by this new vector as compared to the AAV-CMV vector (Figure 5h-j). When this cardiac specific DreAM-plus vector was applied to OsCas12f1-based gene editing, OsCas12f1 protein levels and gene editing outcomes were comparable to the AAV-CMV vector in the heart while being ∼99% eliminated in the liver (Figure 5k-l).

### DreAM-plus reduces the off-target effect of gene editing

The off-target effects of gene editing could be reduced by harnessing delivery methods that transiently express the editors^19,14,26^. However, in situations when such methods are not accessible, whether RNA switch like DreAM-plus could reduce off-target editing in long-lasting vectors remains poorly explored. To answer this question, sgRNAs with validated off-targeted sites were selected for SpCas9^43^, SaCas9^44^ and OsCas12f1^45^. DreAM-plus-controlled vectors were compared to conventional positive control (PC) vectors in the forms of plasmid, lentivirus and AAV to assess off-target effects by amplicon sequencing (Figure 6).

**Fig. 6.**
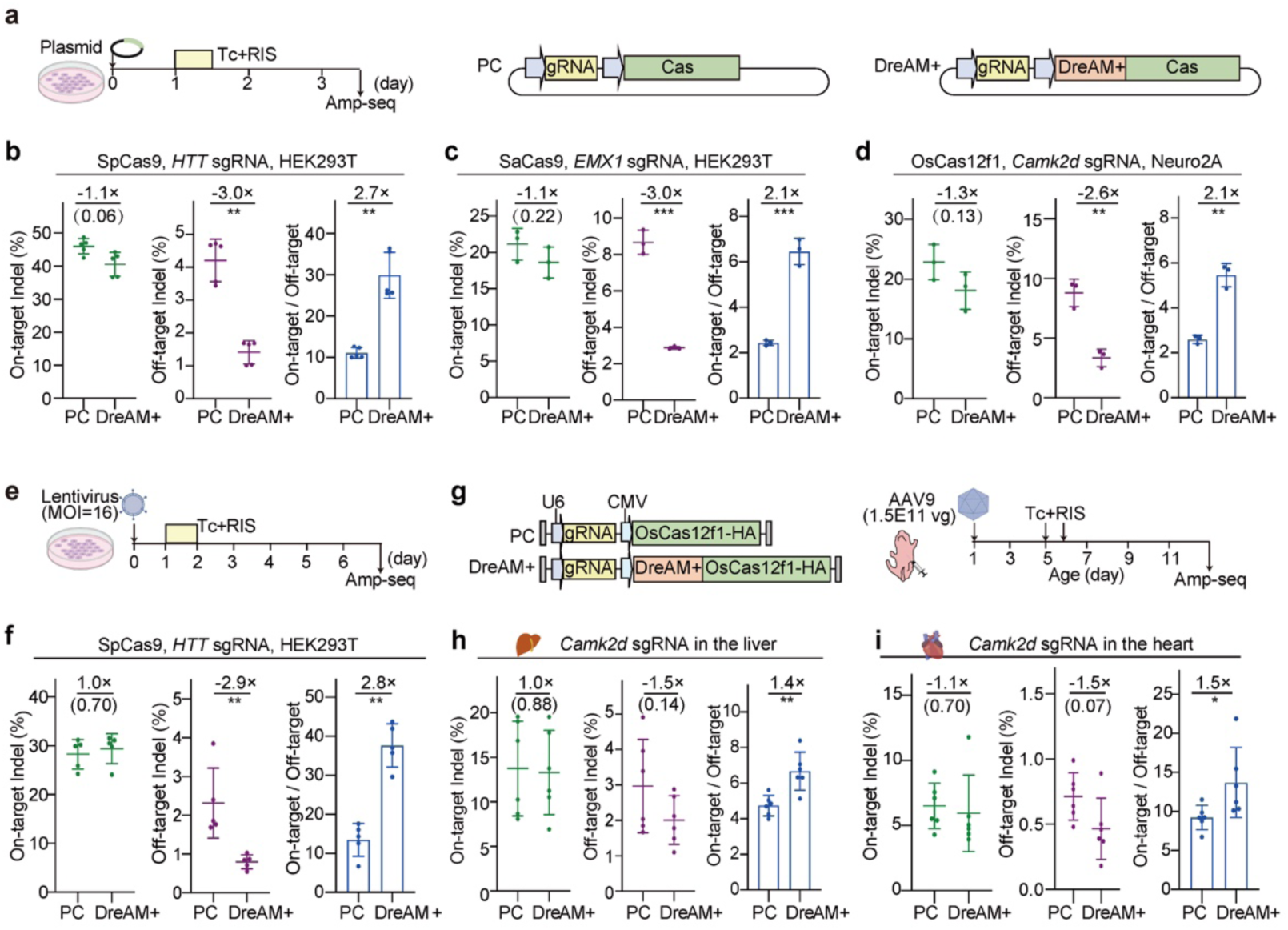
DreAM-plus reduces off-target effects across multiple delivery platforms. **a,** A diagram showing the experimental design to test the off-target effects of PC and DreAM+ plasmids. Cells were transfected for 24 hours before being treated with 0.9 μM tetracycline plus 1.2 μM risdiplam for 12 hours. Then inducers were removed and cells were cultured for an additional 2 days. **b-d,** Analysis of relative editing efficiency at on-target and off-target loci for SpCas9/*HTT*-sgRNA **(b)**, SaCas9/*EMX1*-sgRNA **(c)** and OsCas12f1/*Camk2d*-sgRNA **(d). e,** A diagram showing the experimental design to test the off-target effects of PC and DreAM+ lentivirus. HEK293T cells were transduced with the vectors at a MOI of 16, transiently treated with inducers for 24 hours, and then cultured for 5 days. MOI, multiplicity of infection. **f,** Analysis of relative editing efficiency at on-target and off-target loci for SpCas9/*HTT*-sgRNA lentivirus. **g,** A diagram showing the experimental design to test the off-target effects of PC and DreAM+ AAV. P1 mice were administered with AAV for 4 days before two doses of inducers were applied. One week later, tissues were collected to analyze the off-target effect. **h,i,** Analysis of relative editing efficiency at on-target and off-target loci for OsCas12f1/*Camk2d-*sgRNA in the liver **(h)** and the heart **(i)**. **b-d,f,h,I,** n = 3-6 biological replicates per group. Two-tailed unpaired Student’s t-test: *P < 0.05, **P < 0.01, ***P < 0.001. Non-significant P values are in parentheses.

Through plasmid transfection in HEK293T and Neuro2A cells, a 12-hour treatment of the inducers was sufficient to trigger DreAM-plus-induced gene editing that is comparable to PC at the on-target sites for all three editors (Figure 6a-d). Remarkably, DreAM-plus exhibited 2.6∼3.0-fold reduction of off-target editing while a 2.1∼2.7-fold higher on-target-to-off-target ratio was detected (Figure 6a-d). Similarly, lentivirus-delivered gene editing showed a ∼2.8-fold increase of specificity when the inducers activated DreAM-plus for only 24h (Figure 6e-f). For AAV-delivered in vivo gene editing, two doses of the inducers were injected within 24h before indels were measured a week later (Figure 6g). In both the liver and the heart, the DreAM-plus vector yielded a 1.5-fold higher on-target-to-off-target ratios than the PC vector while the on-target editing remained unaltered (Figure 6h-i). Together, these results demonstrated that DreAM-plus enhanced the specificity of gene editing.

### DreAM-plus reduces the immune toxicity of gene editing

Accumulative studies indicated Cas proteins as robust immunogens that could trigger adaptive immune responses, T-cell-based elimination of host cells, and compromised gene therapy in vivo ^27,29,46^. This concern is further raised as prevalent pre-existing immunity against Cas proteins were found in human population^28,47,48^. To assess if DreAM-plus-mediated transient gene expression could solve this problem, AAV-DreAM-plus-OsCas12f1 and PC vectors were administered to mice before the serum antibody against OsCas12f1 was assessed by the enzyme-linked immunosorbent assay (ELISA) using purified OsCas12f1 proteins as antigen (ED Figure 5a-c). By a month after AAV injection, the DreAM-plus group exhibited significantly reduced OsCas12f1 antibodies than the PC group (ED Figure 5c). However, no significant changes of antigen-induced INFγ-positive cells could be detected among white blood cells or splenocytes by the enzyme-linked immunospot assay (ELISpot) (ED Figure 5d-e). Additionally, flow cytometry failed to identify changed CD8⁺ T cells in the liver and the heart (ED Figure 5f-g).

Next, mouse models with pre-existing immunity against OsCas12f1 proteins were established by subcutaneously injecting purified Cas proteins or LNP with Cas mRNA (Figure 7a). The effective LNP expression was validated in cell culture (ED Figure 6a-b) before administration in mice. ELISpot detected robust antigen-induced INFγ response among splenocytes in the LNP group while Cas proteins elicited less than 10% responses than LNP (Figure 7b). This effect of LNP was also validated by delivering mRNA expressing SaCas9 or Un1Cas12f1 (ED Figure 6c-e). By contrast, Cas protein treatment resulted in five-fold higher titers of anti-Cas antibody in serum than LNP (Figure 7c). Therefore, LNP immunization predominantly elicited a cellular response, while protein immunization primarily induced a humoral response.

**Fig. 7.**
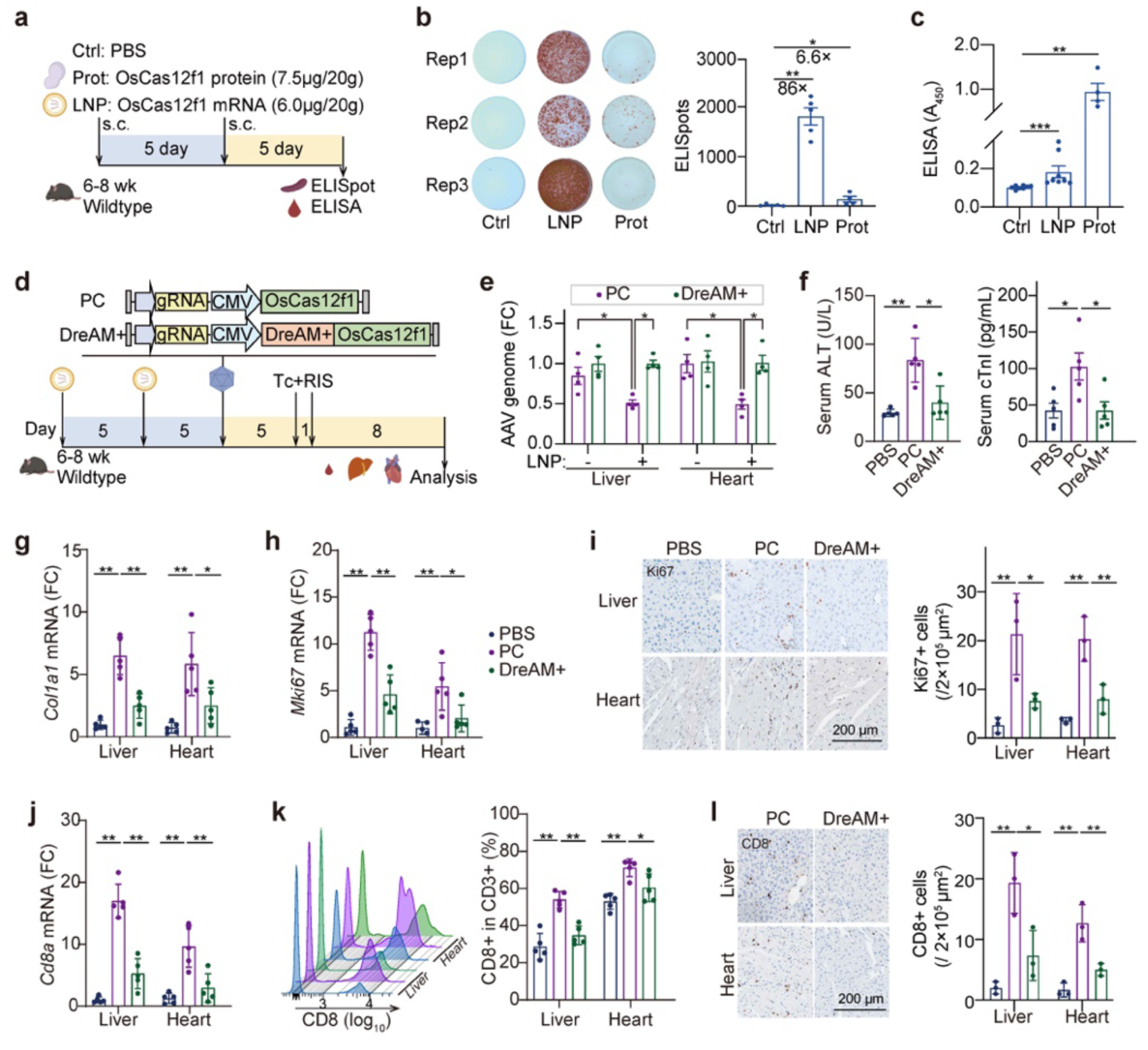
DreAM-plus mitigates cellular immunity against AAV-delivered OsCas12f1 with the presence of LNP-induced pre-existing immunity. **a,** A diagram to establish and assess pre-existing immunity against OsCas12f1 in mice. Prot, protein. LNP, lipid nanoparticle. s.c, subcutaneous injection. **b,** Representative images and quantification of IFNγ-positive ELISpots in splenocytes responding to OsCas12f1 antigen. n=4-5 animals per group. **c,** Quantification of serum anti-OsCas12f1 antibodies by ELISA. n=4 or 8 animals per group. **d,** A diagram to investigate the impact of PC and DreAM+ AAV vectors under pre-existing immunity. Mice were immunized with two doses of LNP-OsCas12f1 before receiving AAV vectors. 80 mg/kg tetracycline plus 50 mg/kg risdiplam were applied daily for two consecutive days. After 8 days, tissues and bloods were collected for analysis. **e,** Quantification of relative AAV genome copies in liver and heart tissues. **f,** Quantification of tissue damage markers alanine transaminase (ALT, left) and cardiac troponin I (cTnI, right) in the serum. **g-h,** mRNA quantification in the liver and heart. **i,** Ki67 immunohistochemistry and quantification. **j-l,** Analysis of CD8⁺ T cell responses via RT-qPCR **(j),** flow cytometry **(k)**, and immunohistochemistry **(l)**. **i,l,** n=3 animals per group. One-way ANOVA with Tukey’s test: *P < 0.05, **P < 0.01. **e-h,j-k,** n=5 animals per group. Mann-Whitney U test: *P < 0.05, **P < 0.01.

The impact of DreAM-plus on AAV-mediated cellular immune response against Cas was then evaluated in the presence of LNP-mediated pre-existing immunity. (Figure 7d). LNP immunization significantly reduced the amount of PC AAV genome in both liver and heart by 50% (Figure 7e), which was an expected result of immune clearance of AAV-transduced cells^28^. Strikingly, DreAM-plus-mediated transient AAV expression completely prevented this effect (Figure 7e). Similarly, serum biomarkers for hepatic (ALT) and cardiac (cTnI) damages were elevated by PC AAV and prevented by DreAM-plus (Figure 7f). DreAM-plus markedly reduced the expression of tissue damage-related markers *Col1a1* and *Mki67* by RT-qPCR, which were validated by immunohistochemistry (Figure 7g-i). RT-qPCR, flow cytometry and immunohistochemistry detected robust CD8-positive T cell infiltration in both the liver and heart, which were attenuated by 60% with DreAM-plus (Figure 7j-l). Noticeably, the effect of DreAM-plus to reduce Cas-associated T-cell cytotoxicity was validated for both OsCas12f1 and UniCas12f1 (ED Figure 7), indicating a generalizable effect.

## Discussion

Precise spatiotemporal control of dynamic gene expression is important in advancing gene function analysis and the outcome of gene therapy. For example, the side effects of gene editing could be mitigated via transient and pulsive expression by electroporation or LNP. However, many tissues are poorly accessibly by these transient delivery approaches, leaving RNA switch-based transient expression by a long-standing vector (such as lentivirus and AAV) an essential alternative.

Recently, several RNA switches have been developed via drug-inducible regulation of a single mechanism, leaving inevitable technical caveats. For instance, this study demonstrated the extensive leaky expression by the aptamer-based Y387 switch and the aberrant expression of dORF-coding proteins by the spliceosome-based Zf2 switch. These problems are challenging to overcome by further optimizing their sequences. Alternatively, this study highlights the integration of multiple miniature RNA switches with distinct mechanisms as a promising solution.

In addition to the reduced leaky expression and the enhanced inducible fold change of DreAM-plus, this new RNA switch exhibited advantages in temporal and spatial parameters. Because tetracycline exhibited shorter half-life than risdiplam, DreAM-plus achieved a minimal transgene duration of less than 24 hours in vivo, which is more transient than the 2∼3-day resolution of DreAM. By combining tissue-specific vectors with administration routes, DreAM-plus allowed tunable gene expression with high specificity in extrahepatic organs like the heart.

The Cas proteins are highly immunogenic and are derived from environmental microbiomes that deposit prevalent pre-existing immunity in human. However, the immunotoxicity of the Cas proteins was poorly studied due to the absence of a robust mouse model. This study harnessed LNP-based vaccination to establish pre-existing cellular immunity against Cas proteins and successfully demonstrated the immunotoxicity of long-term Cas expression by AAV in both the liver and the heart.

Based on this new model, this study showed that DreAM-plus-based transient gene expression as a viable approach to mitigate CD8^+^ T cell infiltration and the immune clearance of cells harboring AAV-Cas.

AAV-based gene editing often requires a dual AAV system due to the large size of the editors. Although RNA switches exhibited smaller sizes than other gene switches, their still exert pressure on the limited payload of AAV vectors^8,31,49^. To overcome this problem, this study successfully adopted the miniature Cas12f editors and achieved switchable gene editing in an all-in-one vector^50^. However, miniature gene editors generally exhibit much lower editing efficiency than canonical editors, demanding further technical improvements before they can be utilized for therapy.

Another limitation of DreAM-plus involves complications with the requirement for two small-molecule inducers. This concern regarding the extra inducer administration could potentially be solved by using safer inducers in the future. For instance, recently the cyclone RNA switch was reported to be induced by acyclovir, which presumably exhibit lower toxicity than tetracycline or risdiplam^11^. In addition, it is intriguing to develop distinct RNA switches on a single inducer. For example, aptamer-based RNA switch could also be developed for risdiplam in the future, which could replace Y387 in DreAM-plus and achieve synergestic multilayer switching by a single inducer.

## Methods

(Additional methods are available in supplementary information)

### Risdiplam and tetracycline

For cell culture, risdiplam powder (T16757, TargetMol, USA) and tetracycline powder (T0912L, TargetMol, USA) were dissolved in DMSO (196055, MPbio) to prepare the 10 mM stock solutions. The stock solutions were aliquoted and stored at −80 °C for up to 1 month before further dilution. For animal studies, tetracycline hydrochloride powder (T0912, TargetMol, USA) and risdiplam powder were dissolved in saline, and the pH was adjusted to 5.

### Animals

Animal experiments were in conformity to the mouse protocol authorized by the Institutional Animal Care and Use Committee of Peking University with the approval number DLASBD0203. C57BL/6 mice were obtained from the Department of Laboratory Animal Science of Peking University Health Science Center and were kept in ventilated cages with appropriate temperature (23 ± 1 °C), humidity (50 ± 5%), illumination (12-h dark–light cycle) and unrestricted access to water and food.

The administration routes of AAVs were determined by animal ages. AAVs were injected to P1 mice subcutaneously. Tail vein injection was performed on adult mice for AAV delivery. Anesthesia was performed via 3% isoflurane (R510-22-10, RWD) inhalation. Animals were euthanized by cervical dislocation following anesthesia.

### Plasmids

Plasmid names and key sequences are listed in Supplementary Table 1-2. In general, DNA fragments were synthesized by Tsingke Biotechnology Co., Ltd. or amplified from pre-existing plasmids. New plasmids were constructed using seamless cloning (B632218, Sangon Biotech).

Plasmids used for transfection and AAV packaging were based on the same backbone. DNA sequences for Y387^7^, Zf2^10^, uP2A, TnpB (addgene#212968), and TnpBmax^51^ were synthesized. DNA fragments for Luciferase (addgene#223188), Un1Cas12f1 (addgene#176544), OsCas12f1 (a gift from Dr Chunlong Xu), AcCas12n (a gift from Dr Quanjiang Ji; addgene#203811) were amplified from pre-existing plasmids. The Tnnt2 promoter of pAAV-Tnnt2-GFP-v2 (addgene#165036) was replaced by the human cytomegalovirus (CMV) promoter to produce an pAAV-CMV-GFP plasmid. Subsequently, Y387, Zf2, Y387- Zf2, Y387- Zf2-uP2A were inserted into pAAV-CMV-GFP plasmid to construct the pAAV-CMV-Y387/ Zf2/ Y387- Zf2/ Y387- Zf2-uP2A-GFP reporter gene plasmids. GFP was replaced by HA-Luciferase to generate pAAV-CMV-Y387/ Zf2/ Y387-Zf2/Y387-Zf2-uP2A-HA-Luciferase. CMV was replaced by Tnnt2 and miR122TS was inserted to generate cardiac-specific vectors. The reporter genes were replaced by Cas or TnpB sequences, and U6-sgRNA or reRNA was inserted to generate plasmids for gene editing. For gene editing, gRNA sequences are summarized in Supplementary Table 3.

For lentivirus plasmid construction, backbone plasmid pLenti-U6-sgRNA-EFS-SpCas9-FLAG-P2A-Puro was bought from Tsingke Biotechnology Co., Ltd. *HTT* sgRNA was inserted into the backbone plasmid to construct the pLenti-U6-*HTT* sgRNA-DreAM+-EFS-SpCas9-flag. DreAM+ was inserted to generate pLenti-U6-*HTT* sgRNA-EFS-SpCas9-flag plasmid.

### RT-qPCR

For mRNA level analysis, total RNA was purified using the TransZol Up Plus RNA Kit (ER501-01-V2, TransGene) with genomic DNA removed. RNA was reverse transcribed with HiScript III All-in-One RT SuperMix (R333-01, Vazyme Biotech). Relative quantitation of gene expression was determined by normalizing to *Gapdh*.

For tissue AAV genome copies analysis, genomic DNA was extracted using TIANamp Genomic DNA Kit (DP304, TIANGEN, China). Relative quantitation of AAV genome copies in the liver and heart was determined by normalizing to *Tnni3a*.

For SYBR qPCR, each sample was assayed in three technical replicates using the QuantStudio 3 Real-Time PCR System (Thermo Fisher Scientific) with 2×Taq Pro Universal SYBR qPCR Master Mix (Q712-02, Vazyme Biotech). Primers for qPCR are listed in Supplementary Table 4.

### Amp-seq analysis

For RNA splicing analysis, the total RNA was extracted, and genomic DNA was removed using DNaseI (RT411, TIANGEN). Then, 1 µg of RNA was reverse transcribed. Subsequently, cDNA was amplified using 2×Taq PCR Mix (KT201, TIANGEN) and amp-seq primers (Supplementary Table 2). The PCR products were purified using the TIANgel Purification Kit (DP214, TIANGEN) before sequencing.

The sequencing libraries were generated and sequenced on Illumina platforms using paired end PE250 reads on an Illumina NovaSeq 6000 platform at Novogene. For the alignment workflow, custom reference genomes (FASTA) and annotation files (GTF) were constructed for the pA-Y387 and DreAM-Zf2 controls, the full-length Y387-DreAM fusion system (containing segments A-E), and the deletion variants (ΔA, ΔC, and ΔAC). Raw sequencing data underwent quality control using FastQC (v0.12.1). Subsequently, low-quality bases and adapters were removed using Trim Galore (v0.6.10). The filtered reads were mapped to their corresponding custom references using STAR (v2.7.11b). The resulting alignment files were processed with Samtools (v1.19.2) for sorting and indexing.

For quantitative analysis, a custom Python pipeline was implemented to determine the Percent Spliced In (PSI) values. By parsing CIGAR (Concise Idiosyncratic Gapped Alignment Report) information from the BAM files, the script identified reads spanning specific exon-exon junctions. The PSI was calculated by quantifying the read coverage across splice junctions characteristic of each isoform, thereby evaluating the necessity of specific sequence fragments for the splicing switch.

For gene editing analysis, genomic DNA was extracted using TIANamp Genomic DNA Kit (DP304, TIANGEN, China). The targeted loci for sequencing were amplified using Taq PCR MasterMix (KT211, TIANGEN, China) and purified by TIANgel Purification Kit (DP219, TIANGEN). Sequencing was performed on an DNBSEQ-T7 platform at Geneplus, China. Sequencing results were analyzed by CRISPResso2^52^. See Supplementary Table 3 for primer sequences for each gRNA.

### Western blot

Cells or tissues were washed with ice-cold phosphate-buffered saline (PBS) before lysis in RIPA buffer (P0012B, Beyotime Biotechnology). After centrifugation, the supernatant was diluted in 4× sodium dodecyl sulphate-polyacrylamide gel electrophoresis (SDS-PAGE) loading buffer (B1016, Solarbio). After being boiled at 70 °C for 10 min, proteins were separated on 4–12% SurePAGE gradient gel (M00725, GenScript), transferred to Immobilon-P PVDF membrane (IPVH00010, Merck Life Science) and blocked by 5% non-fat milk–TBST (Tris buffered saline with 0.1 % Tween 20) at 25 °C for 1 h. Membranes were incubated with the primary antibodies overnight at 4 °C, followed by 3 × 5 min TBST washes. The membranes were probed by horseradish peroxidase-conjugated secondary antibodies for 1 h at room temperature, followed by 3 × 5 min of TBST washes. After adding ECL Western Blotting Substrate (PE0010, Solarbio), chemiluminescence was detected by an iBright CL1500 Imaging System (Thermo Fisher Scientific). Antibodies used in this study are listed in Supplementary Table 5. Uncropped western blots can be found in Source Data.

### ELISpot analysis

Splenocytes were isolated aseptically from sacrificed mice. Briefly, the spleen was homogenized gently through a 70-μm cell strainer using the plunger of a 3-mL syringe. For blood cells analysis, plasma was collected at indicated time point. Erythrocytes were lysed using ACK lysis buffer, and the remaining cells were washed twice with complete RPMI-1640 medium (supplemented with 10% fetal bovine serum, 1% penicillin and streptomycin). Cell viability and concentration were determined by trypan blue exclusion.

Mouse IFNγ ELISPOT Kit (551881, BD) was used for the ELISpot assay. In brief, a 96-well polyvinylidene difluoride (PVDF)-backed plate pre-coated with anti-mouse IFNγ capture antibody was used according to the manufacturer’s protocol. The plate was blocked with complete RPMI-1640 medium for 1 hour at room temperature. Freshly isolated splenocytes were then seeded into the plate at a density of 2.5 × 10⁵ cells per well in triplicate and stimulated with the commercialized specific protein antigen at a final concentration of 2.5 μg/mL. The plate was incubated for 20 hours in a humidified incubator at 37°C with 5% CO₂.

After incubation, cells were removed by washing the plate extensively with PBS containing 0.05% Tween-20 (PBST). The biotinylated anti-mouse IFN-γ detection antibody was added and incubated for 2 hours at room temperature, followed by incubation with Streptavidin-HRP for 1 hour. Spot-forming units (SFUs) were visualized by using the AEC substrate set (551951, BD) developed until distinct spots emerged. The reaction was stopped by rinsing the plate with distilled water. The plate was air-dried in the dark, and spots were counted using an automated ELISPOT reader system (AID).

### ELISA analysis of anti-Cas12f1 antibody

Cas12f1 ELISA protocol was adapted from Moreno AM et al^29^. In brief, purified OsCas12f1 protein (JS0425, OriGene) or purified Un1Cas12f1 protein (32119, TOLOBIO) was diluted with coating buffer (421701, Biolegend). MaxiSorp ELISA plates (423501, Biolegend) then were coated with 0.5 μg/100 μL Cas12f1 protein per well and incubated at 4°C overnight. The plates were washed 3 times for 5 min with 200 μL of PBST and subsequently blocked with 200 μL of BSA blocking solution at room temperature for 2 h. The washing was repeated again. Serum samples were added at 1:5 dilution and the plates were incubated at 37°C for 4 h. The wells were washed 3 times for 5 min each and 100 μL of HRP-labeled goat anti-mouse IgG1, which was diluted with BSA blocking solution at a ratio of 1:5000, was added to each well. After 1 h incubation at room temperature, the wells were washed 4 times for 5 min each and then 100 μL of 3,3′,5,5′-tetramethylbenzidine (TMB) substrate (421101, Biolegend) was added to each well and incubated for 15 min at room temperature in the dark. The optical density (OD450) at 450 nm was measured using CLARIOstar Plus multi-mode microplate reader after adding 100 μL of stop solution to each well.

### Statistical analysis

Statistical analysis and plotting were performed using GraphPad Prism 10. Statistical tests are described in figure legends. The following values were statistically significant: *p < 0.05, **p < 0.01, ***p< 0.001, ****p< 0.0001. Data are presented using the mean ± s.d. in bar plots. Dots represent raw data points.

## Contributions

Conceptualization, Y.G., Yueyang Zhang, L.M., F.G.;

Experimental studies, Yueyang Zhang, Z.L., Y.L., G.C., Z.Z., Y.X.;

Data collection and analysis, Yueyang Zhang, Y.Y., D.Z., T.L., Yaohua Zhang;

Supervision, Y.G., L.M., F.G. and K.Y.;

Writing, Y.G. and Yueyang Zhang.

## Acknowledgements

This study was funded by Beijing Natural Science Foundation (F252059 to Y.G. and F.G.), National Natural Science Foundation of China (82570307 to Y.G.), the Science and Technology Plan Project of Tongzhou District, Beijing (WS2025014 to F.G.) and Vituner Therapeutics.

## Data availability

Sequencing data are available from the Genome Sequence Archive (https://ngdc.cncb.ac.cn/gsa/, CRA037381). Source data are provided with this paper.

---

**Extended Data Fig.1.**
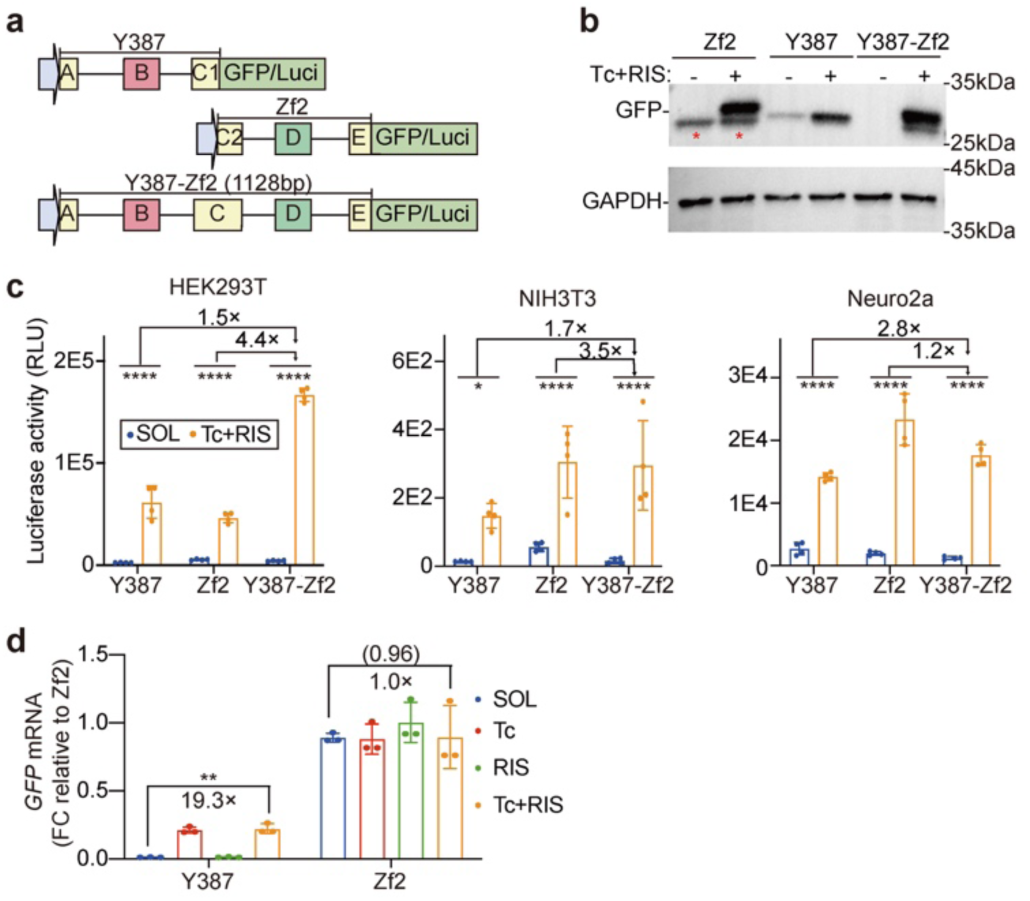
Comparative analysis of the Y387-Zf2 fusion element versus individual Y387 and Zf2 switches in vitro. **a,** A diagram showing reporter vector constructs. Luci, Firefly luciferase. **b,** Western blot analysis of GFP controlled by the indicated RNA switch in HEK293T cells treated for 24 hours with solvent or 1 µM tetracycline plus 1 µM risdiplam. The red stars indicate N-terminal truncated peptides by dORF. **c,** Quantification of luciferase activity under the control of indicated switches. The fold changes of induced dynamic ranges were labeled above. RLU, relative light unit. n=4 biological repeats. Two-tailed unpaired Student’s t-test: *P < 0.05, **P < 0.01, ****P < 0.0001. **d,** *GFP* mRNA levels in HEK293T cells with indicated switches and inducers. n=3 biological repeats. One-way ANOVA with Tukey’s test: **P < 0.01. Non-significant P values are in parentheses.

**Extended Data Fig.2.**
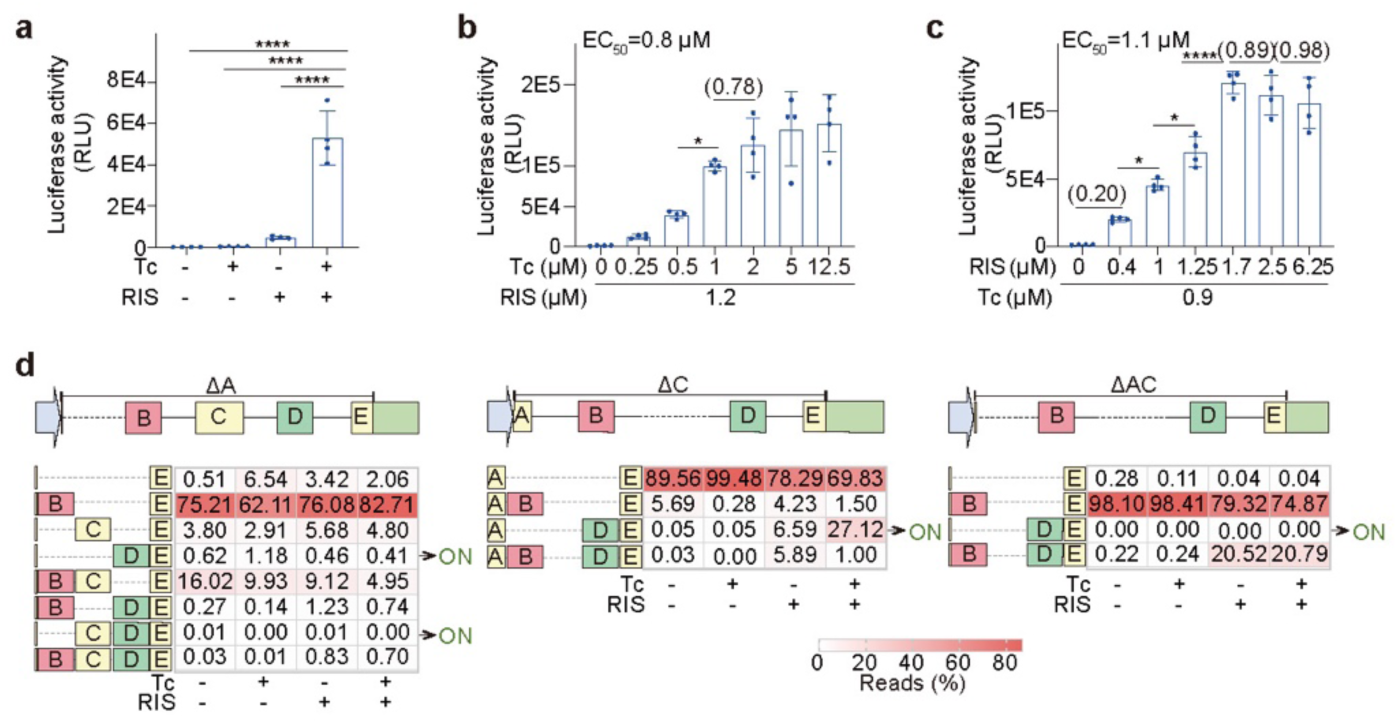
In vitro characterization of Y387-Zf2 and its truncated variants. **a,** Quantification of luciferase activity under the control of Y387-Zf2 in HEK293T cells treated with solvent, 1 μM tetracycline, 1 μM risdiplam or both for 24 hours. **b,c,** Quantification of luciferase activity in response to a gradient of tetracycline or risdiplam for 24 hours. **d,** Heatmap showing the average ratio of splicing variants generated by different truncated switches. n=3 biological repeats. ON indicates splicing variants with exon D but without exon B. **a-c,** n=4 biological repeats. One-way ANOVA with Tukey’s test: *P < 0.05, ****P < 0.0001. Non-significant P values are in parentheses.

**Extended Data Fig.3.**
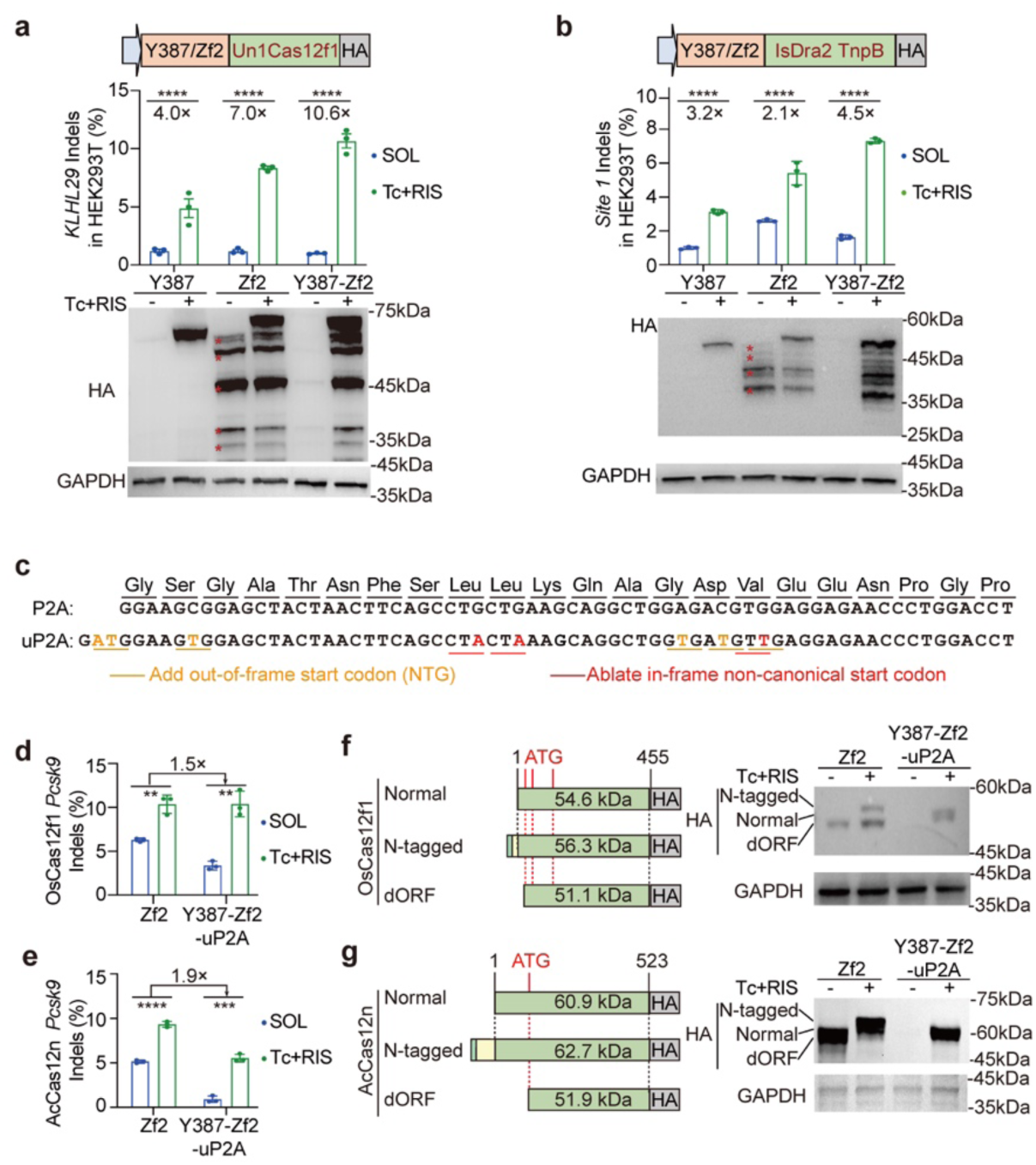
DreAM-plus is broadly applicable for gene editing. **a,b,** Editing efficiency and western blot analysis of Un1Cas12f1 **(a)** and IsDra2 TnpB-**(b)**. Performance of each editor was evaluated by treatment with solvent or 0.9 μM tetracycline plus 1.2 μM risdiplam for 48 hours. **c,** DNA and amino acid sequences of P2A and uP2A. Out-of-frame start codons (orange underscored) were introduced into uP2A and in-frame non-canonical start codons (red underscored) were ablated. **d,e,** Gene editing analysis of OsCas12f1 **(d)** and AcCas12n **(e)** in N2a cells. **f,g,** A diagram showing the predicted molecular weights of proteins translated at distinct start codons and the corresponding western blot bands of OsCas12f1 **(f)** and AcCas12n **(g)**. **a,b,d,e,** n = 3 biological repeats. Two-tailed unpaired Student’s t-test: **P < 0.01, ***P < 0.001, ****P < 0.0001.

**Extended Data Fig.4.**
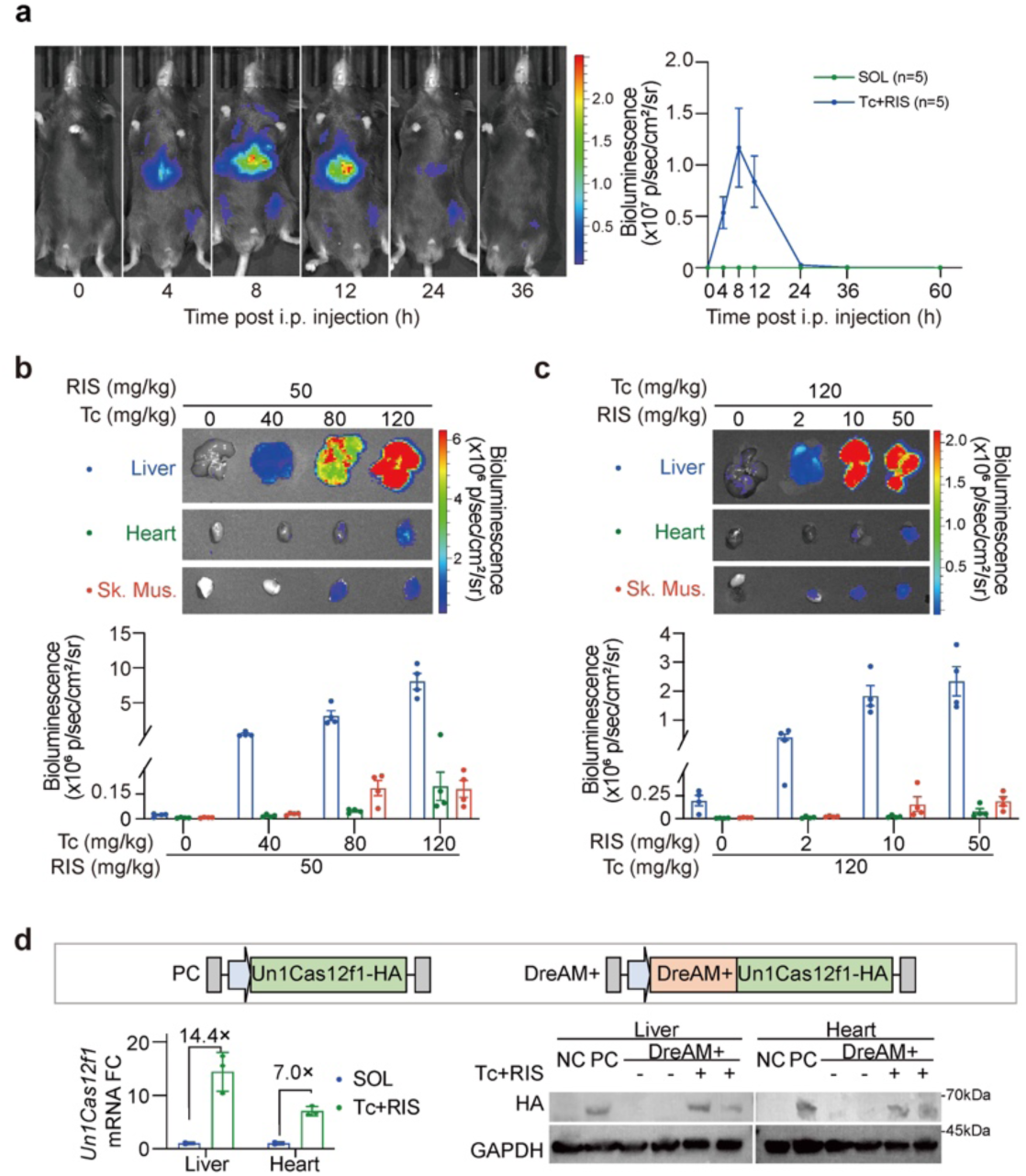
DreAM-plus mediates in vivo regulation across multiple organs. **a,** Bioluminescence images and quantification of luciferase signals at the indicated time points after administration of 80 mg/kg tetracycline plus 50 mg/kg risdiplam. i.p., intraperitoneal injection. n = 5 animals per group. **b,c,** Bioluminescence images and quantification of signals from isolated organs. Mice were treated with a gradient of tetracycline **(b)** or risdiplam **(c)** concentrations for 12 hours. Sk. Mus., skeletal muscle. n = 4 animals per group. **d,** RT-qPCR and western blot analysis of Un1Cas12f1 in the liver and heart. P1 mice were administered with 1.5E12vg AAV9 via subcutaneous injection. At P6 and P7, mice were treated with 2 doses of solvent or inducers. Tissues were harvested at P8. n = 3 animals per group.

**Extended Data Fig.5.**
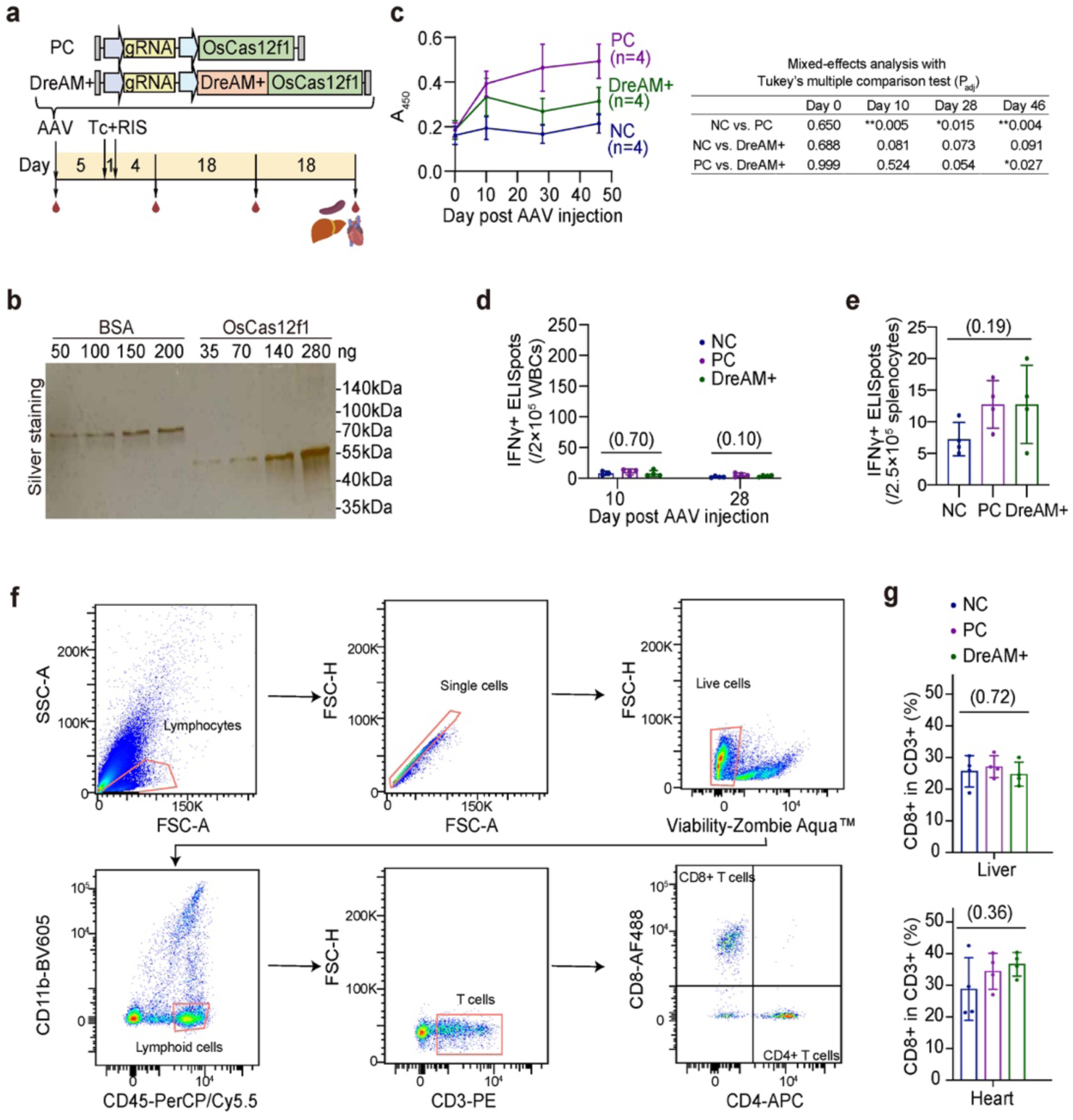
DreAM-plus mitigates the humoral immune response against AAV-delivered OsCas12f1. **a,** Experimental design to evaluate immune responses to AAV-delivered OsCas12f1 in mice. **b,** Validation of protein purification by silver stain. BSA was used as standards for quantification. BSA, bovine serum albumin. **c,** ELISA analysis of serum anti-OsCas12f1 antibody at indicated time points. n=4 animals per group. NC, PBS-treated negative control. **d,e,** Quantification of IFNγ-positive ELISpots generated by antigen-stimulated white blood cells (WBCs)**(d)** or splenocytes **(e)**. **f,** Gating strategy for CD8+ T cell analysis in tissues. **g,** Quantification of CD8+ cell proportion among CD3+ T cells. **d,e,g,** n = 3-4 animals per group. One-way ANOVA with Tukey’s test, non-significant P values are in parentheses.

**Extended Data Fig.6.**
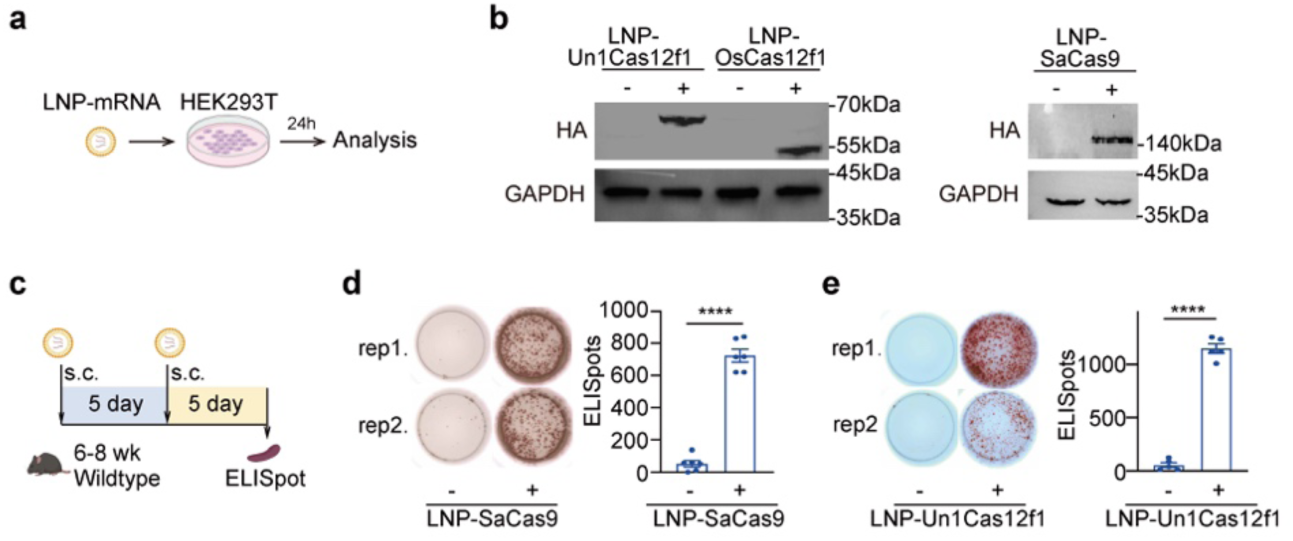
LNP-mRNA triggers pre-existing cellular immunity against Cas. **a,** Experimental design to test LNP-mRNA expression in HEK293T cells. **b,** Western blot analysis of Un1Cas12f1, OsCas12f1 and SaCas9 by LNP-encapsulated mRNAs. **c,** Experimental design to generate LNP-induced pre-existing immunity. **d,e,** Images and quantifications of antigen-induced IFNγ-positive ELISpots in splenocytes for LNP-SaCas9 **(d)** or LNP-Un1Cas12f1 **(e)**. n = 5 or 6 animals per group. Two-tailed unpaired Student’s t-test: ****P < 0.0001.

**Extended Data Fig.7.**
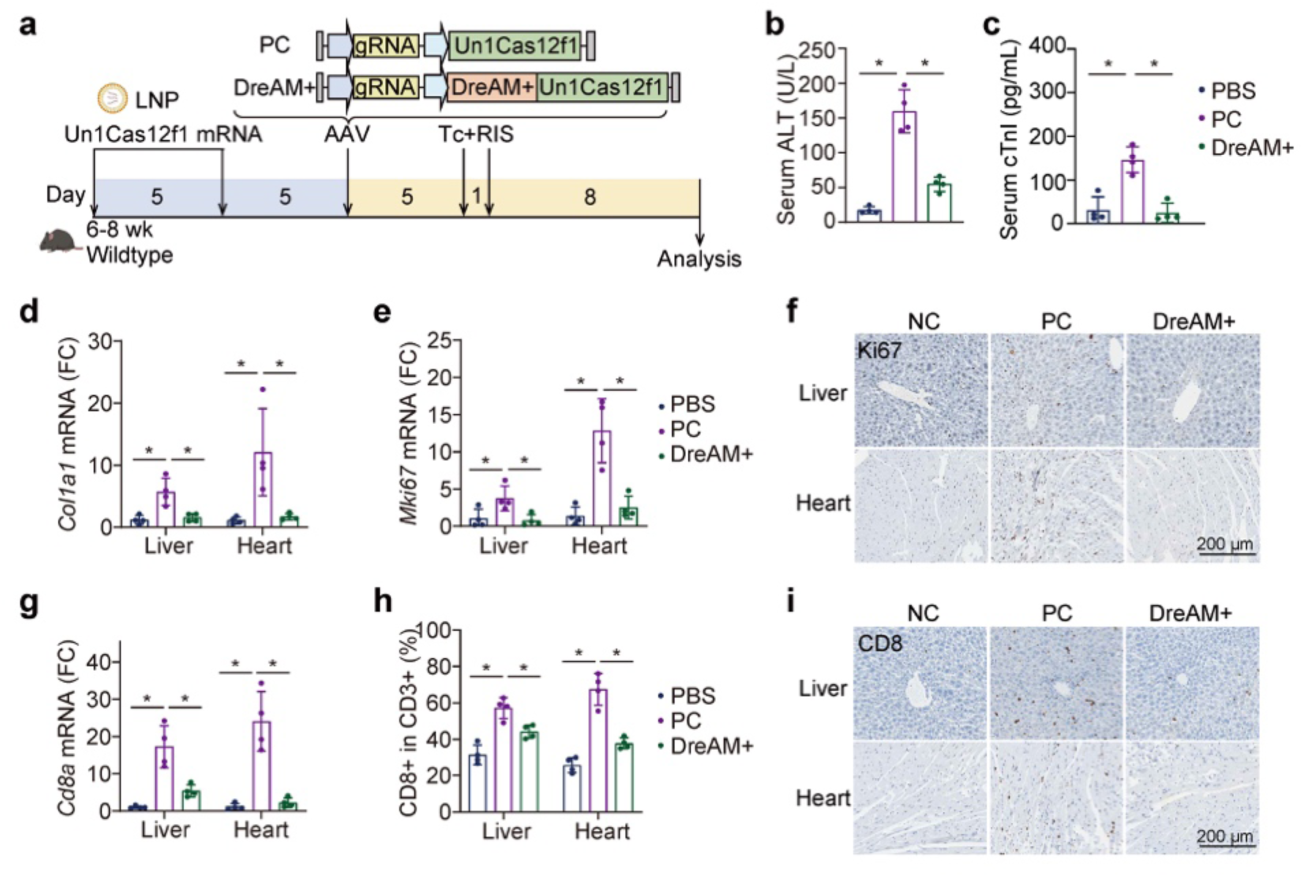
DreAM-plus mitigates the immune response against Un1Cas12f1 with pre-existing immunity. **a,** Experimental design to investigate the impact of DreAM+ on Un1Cas12f1 immunity at the presence of pre-existing immunity. **b,c,** Quantification of liver damage marker ALT **(b)** and heart damage marker cTnI **(c)** in the serum. **d,** *Col1a1* mRNA quantification. **e,f,** *Mki67* mRNA quantification **(e)** and immunohistochemistry analysis **(f)**. **g-i,** Analysis of CD8⁺ T cell infiltration in the liver and heart by *CD8a* mRNA qPCR **(g),** CD8⁺ cell flow cytometry **(h)** and immunohistochemistry analysis of CD8 **(i)**. **b-e,g,h** n=4 animals per group. Mann-Whitney U test: *P < 0.05.

