## supplementary methods for "Synergistic integration of inducible RNA switches enhances the manipulation of vector expression"

Zhang et al.

Supplementary Information

### Supplementary Methods

Cell culture, fluorescence analysis and in vitro luciferase reporter gene assay

HEK293T (CL-0005), NIH3T3 (CL-0171), and Neuro2a (CL-0168) cells were obtained from Procell, China. These cells were cultured in high-glucose DMEM (HK2109.07, HUANKE) containing 10% Tetracycline-free FBS (C2720, Vivacell) and 1× penicillin–streptomycin solution (BC-CE-007, Bio-channel) at 37 °C and 5% CO2 with saturating humidity. Cells were plated at a density of $1.5 \times 10^5$ cells per well in 24-well plates for 12 h before plasmid transfection or virus transduction.

Lipo8000 (C0533, Beyotime, China) was used following the manufacturer's instructions during plasmid transfection. Lentivirus transduction was performed with 0.1 % polybrene (40804ES76, Yeasen). For plasmid transfection, the inducers or solvent and transfection complexes were added to cells simultaneously. For virus transduction, cells were treated with fresh medium containing the inducers or solvent after 24 h of virus transduction.

The fluorescence intensity of cultured cells was measured on a CLARIOstar Plus multi-mode microplate reader (BMG LABTEC) at room temperature. Additionally, images were taken under a fluorescence microscope (BZ-X810, KEYENCE) and images were quantified by ImageJ (version 1.54 f).

Luciferase activity was determined using a luciferase reporter gene assay kit (11401ES, Yeason). Briefly, cells were seeded in 96-well plates and allowed to adhere overnight. Subsequently, cells were transfected with the firefly luciferase reporter plasmid using Lipo8000 and treated with tetracycline or risdiplam at the indicated concentrations. After 24 hours, the culture medium was aspirated, and cells were washed gently with 1×PBS. 100 μL lysis buffer (provided in the kit) was added to each well. For the luciferase activity measurement, 20 μL of the cell lysates was aliquoted into a white opaque 96-well assay plate. The firefly luciferase activity was measured immediately by injecting 100 μL of Luciferase Assay Substrate into each well using CLARIOstar Plus multi-mode microplate reader with luminescence detection capability.

Viral vector production

AAVs were produced by PackGene Biotech. In brief, 140 μg AAV-ITR, 140 μg AAV-Rep/Cap and 320 μg pHelper (pAd-deltaF6, Penn Vector Core) plasmids were produced using Qiagen Plasmid Kits (12165, Qiagen) and transfected into 10 15-cm plates of HEK293T cells using polyethylenimine (PEI) transfection reagent (40816ES03, Yeasen). Then, 60 h after transfection, cells were scraped out, resuspended in PBS and lysed by 4 freeze–thaw–vortex cycles. Cell debris was removed after 10,000 g centrifugation. AAVs in cell culture medium were precipitated by 8% PEG8000 (P2139, Sigma-Aldrich), resuspended in PBS and pooled with cell lysates. AAVs were purified in a density gradient (D1556-250ML, Sigma) by ultracentrifugation and concentrated in PBS using an ultrafiltration tube (100K molecular weight cutoff, Millipore). AAV titer was absolutely quantified by qPCR using primers amplifying a fragment of inverted terminal repeat (ITR).

Lentivirus production was performed by PackGene Biotech. In brief, psPAX2, pMD2.G and pLenti-U6-HTT-sgRNA-DreAM+-EFS-SpCas9-flag or pLenti-U6-HTT-sgRNA-EFS-SpCas9-flag were produced using Qiagen Plasmid Kits (12165, Qiagen). Triple transfection was performed via the PEI method into 10-cm plates of HEK293T cells. Viruses were collected at 48 h, 72 h and 96 h post transfection from the cell culture medium after centrifugation at 880 g for 5 min. The lentiviruses were then purified through ultracentrifugation and concentrated using an ultrafiltration tube. The lentivirus titer was absolutely quantified by qPCR using primers amplifying a fragment of long terminal repeat (LTR).

mRNA synthesis, LNP production and characterization

mRNAs were synthesized by in vitro transcription from plasmids containing T7 promoter, CDS region, 5' and 3' untranslated regions (UTRs) and a poly A tail (~100 nt). In vitro transcription was performed using EasyCap T7 Co-transcription Kit with CAG Trimer Kit (DD4203-01, Vazyme). The synthesized modRNAs were purified by RNA clean beads (N412-01, Vazyme). The T7-5’UTR-CDS-3’UTR-polyA sequences of mRNAs used in the current study are listed in Supplementary Table 1.

Lipid nanoparticles (LNPs) were prepared using microfluidic technology by mixing an aqueous phase containing mRNA in citrate buffer (pH=4) with an ethanol solution containing four lipid components at a volume ratio of 3:1. The molar ratio of the lipids in the ethanol solution was ALC-0315:DSPC:cholesterol:DMG-PEG2000 = 50:10:38.5:1.5. After preparation, the LNPs were transferred into a 10 kDa dialysis bag and dialyzed overnight against 1× PBS at 4°C. The mRNA concentration and encapsulation efficiency were measured using the Quant-iT RiboGreen RNA assay. The hydrodynamic diameter and zeta potential of the LNPs were determined via dynamic light scattering after 100-fold dilution in either 1× PBS or 4 mM KCl.

Tissue sectioning and immunohistochemistry analysis

Immunohistochemistry (IHC) was performed by Wuhan Servicebio Technology. Formalin-fixed, paraffin-embedded (FFPE) tissue blocks were sectioned at a thickness of 4-5 μm using a microtome. Prior to staining, tissue sections were deparaffinized in xylene (twice, 10 minutes each) and rehydrated through a graded ethanol series (100%, 95%, 80%, 70%; 5 minutes each), followed by a final rinse in distilled water.

For antigen retrieval, slides were immersed in pre-heated citrate-based antigen retrieval buffer (pH 6.0) or EDTA buffer (pH 9.0) and heated in a microwave oven or a pressure cooker for 15-20 minutes. Endogenous peroxidase activity was quenched by incubating sections with 3% hydrogen peroxide in methanol for 15 minutes at room temperature in the dark, followed by three PBS washes. Non-specific binding sites were blocked by applying 5% normal goat serum (or serum matching the host species of the secondary antibody) in PBS for 1 hour at room temperature.

Sections were incubated overnight at 4°C with the primary antibody. Antibodies used in immunohistochemistry are listed in Supplementary Table 5. After three washes in PBS, sections were incubated with a horseradish peroxidase (HRP)-conjugated secondary antibody for 1 hour at room temperature. Following another three PBS washes, the peroxidase signal was developed using a 3,3'-diaminobenzidine (DAB) chromogen substrate kit. The reaction was stopped by immersing slides in distilled water. Sections were counterstained with Mayer's hematoxylin for 1-2 minutes, followed by differentiation in 1% acid alcohol and bluing in tap water. Subsequently, sections were dehydrated through an ascending ethanol series (70%, 80%, 95%, 100%), cleared in xylene, and mounted with a permanent mounting medium under a coverslip. Stained slides were scanned using OCUS 40 microscope slide scanners (Grundium). Ki-67 or CD8 positive cells were counted and quantified.

Bioluminescence imaging

Sterile D-PBS (60152ES76, Yeasen) and D-Luciferin (40902ES03, Yeasen) were used to prepare 15 mg ml−1 luciferin solution, which were injected intraperitoneally 10 min before imaging at the luciferin to body weight ratio of 150 mg kg−1. Animals were anesthetized via 3% isoflurane inhalation before bioluminescent imaging and analysis in an IVIS Spectrum In Vivo Imaging System (PerkinElmer).

#### Flow cytometry

Mice were perfused with 20-30 mL of ice-cold, sterile PBS to remove circulating blood cells from the organs. The heart and liver were then aseptically excised, weighed, and placed in separate petri dishes containing cold PBS. For Heart tissue digestion, the ventricle was minced into fine pieces and digested in 1 mL of digestion medium (RPMI-1640 containing 1 mg/mL collagenase I, 0.1 mg/mL DNase I) at 37°C for 45 minutes with gentle agitation. For liver tissue digestion, the liver lobe was minced into fine pieces and digested in 1 mL of digestion medium (RPMI-1640 containing 1 mg/mL collagenase IV, 0.1 mg/mL DNase I,) at 37°C for 30 minutes with gentle agitation. The digested tissue was passed through a 70-μm cell strainer. The filtrate was centrifuged at 500 × g for 5 minutes at 4°C. Erythrocytes were lysed using ACK lysis buffer (102420, Yimi Biotech). Cells were incubated with an Fc receptor blocking antibody (anti-mouse CD16/32) for 10 minutes on ice. Subsequently, cells were stained with viability dye, followed by a cocktail of fluorochrome-conjugated antibodies against cell surface markers for 30 minutes in the dark at 4°C. After staining, cells were washed twice with FACS buffer. Data was acquired on a BD FACSymphony™ A1 cell analyzer and analyzed using Flowjo 10.10.0 software. Antibodies and dye used in flow cytometry are listed in Supplementary Table 5.

#### Blood test

Blood was allowed to clot for 1 hour at room temperature. After centrifugation at 4,000 rpm for 10 min, serum was obtained from the supernatant. Serum components were measured using ALT assay kit (C009-2-1, Nanjing Jiancheng Bioengineering Institute), mouse TNNI3/cTn-I ELISA kit (E-EL-M1203, Elabscience) following the manufacturer's instructions.
